# Elucidating the metabolic roles of isoflavone synthase-mediated protein-protein interactions in yeast

**DOI:** 10.1101/2024.10.24.620109

**Authors:** Chang Liu, Jianing Han, Sijin Li

## Abstract

Transient plant enzyme complexes formed via protein-protein interactions (PPIs) play crucial regulatory roles in secondary metabolism. Complexes assembled on cytochrome P450s (CYPs) are challenging to characterize metabolically due to difficulties in decoupling the PPIs’ metabolic impacts from the CYPs’ catalytic activities. Here, we developed a yeast-based synthetic biology approach to elucidate the metabolic roles of PPIs between a soybean-derived CYP, isoflavone synthase (GmIFS2), and other enzymes in isoflavonoid metabolism. By reconstructing multiple complex variants with an inactive GmIFS2 in yeast, we found that GmIFS2-mediated PPIs can regulate metabolic flux between two competing pathways producing deoxyisoflavonoids and isoflavonoids. Specifically, GmIFS2 can recruit chalcone synthase (GmCHS7) and chalcone reductase (GmCHR5) to enhance deoxyisoflavonoid production or GmCHS7 and chalcone isomerase (GmCHI1B1) to enhance isoflavonoid production. Additionally, we identified and characterized two novel isoflavone *O*-methyltransferases interacting with GmIFS2. This study highlights the potential of yeast synthetic biology for characterizing CYP-mediated complexes.

## Main

Plant enzyme complexes, sometimes called metabolons, are supramolecular complexes of sequential metabolic enzymes in a pathway formed through transient protein-protein interactions (PPIs)^1^. These complexes are considered important regulatory machinery in plant natural product (PNP) biosynthesis with diverse metabolic functions. Multiple recent studies have shown that enzyme complexes assembled on a non-catalytic scaffold protein can provide rapid and reversible control of metabolic flux, thereby enhancing PNP production specificity or efficiency. For example, the PPI between the non-catalytic chalcone isomerase-like protein (CHIL) and chalcone synthase (CHS), which have been found across various land plants, enhances CHS’s catalytic specificity to favor the production of chalcone over byproducts such as p-coumaroyltriacetic acid lactone^2^. Another example involves membrane steroid-binding proteins (MSBPs), which organize monolignol cytochrome P450 (CYP) enzymes to increase lignin production in *Arabidopsis thaliana*^3^. However, many plant enzyme complexes use functional enzymes rather than non-catalytic proteins as scaffolds and remain challenging to characterize. These scaffold enzymes, usually CYPs, anchor the complex to organelles such as endoplasmic reticulum (ER), thereby playing dual roles as both catalysts and modulators that regulate the activities of other enzymes. Examples include the cinnamate 4-hydroxylase (C4H) assembling a multi-enzyme complex in tobacco’s phenylpropanoid metabolism, flavone synthase (FNS) and flavonoid 3’-hydroxylase (F3’H) in the flavonoid biosynthesis of snapdragon and torenia^4^, and three CYPs in *A. thaliana’*s camalexin metabolism^5^. The involvement of the ER membrane between the scaffold CYP and cytosolic, catalytic enzymes poses unique challenges for in vitro functional characterization. More importantly, it is difficult to generate a non-catalytic variant of the scaffold CYP without interrupting its PPIs with other enzymes for both in planta and in vitro assays.

Baker’s yeast (*Saccharomyces cerevisiae*) provides a feasible approach to reconstitute and characterize the CYP-mediated plant enzyme complexes heterologously. Numerous synthetic biology studies have proven that ER-bound CYPs derived from plants can be functionally expressed and localized correctly into yeast ER^6–8^, paving the way to ER-bound complex characterization. Particularly, the supply of CYP’s essential redox partner, namely cytochrome P450 reductase (CPR), is insufficient in yeast, making it uniquely suitable for CYP-mediated complex characterization. Co-expressing a plant-derived CYP with its interacting enzyme in a CPR-deficient yeast can easily deactivate the CYP while maintaining the interactions. Thus, reconstituting membrane-bound plant enzyme complexes in yeast can be a promising approach to characterize the impact of the PPI on PNP synthesis, ultimately shedding light on new strategies for PNP biomanufacturing and plant engineering.

This work presents a yeast-based synthetic biology method to elucidate the metabolic role of CYP-mediated plant enzyme complexes in vivo. We chose to develop the proposed method on the soybean (*Glycine max*) enzyme complexes involved in isoflavonoid (including both deoxyisoflavonoid and isoflavonoid, referred to as (deoxy)isoflavonoid in this work for clarity) metabolism. (Deoxy)isoflavonoids are an important class of PNPs primarily produced by legumes having potential health promoting effects. Their derivatives such as glyceollins are crucial phytoalexins regulating plant-microbe interactions^9^. There are two competing pathway branches in soybean producing deoxyisoflavonoids (e.g., daidzein^10^) or isoflavonoids (e.g., genistein^11^), respectively, yet how these two pathways are coordinated dynamically and swiftly in planta has not been fully understood. The deoxyisoflavonoid branch starts with concerted reactions catalyzed by chalcone synthase (GmCHS) and chalcone reductase (GmCHR)^12^ toward isoliquiritigenin, which converts to liquiritigenin spontaneously^13,14^ or by chalcone isomerase (GmCHI). Isoflavone synthase (GmIFS^15^), an ER-bound CYP, catalyzes the hydroxylation and aryl-ring migration of liquiritigenin to 2-hydroxyliquiritigenin, which then dehydrates to daidzein spontaneously or by 2-hydroxyisoflavanone dehydratase (GmHID)^16^ . In the isoflavonoid branch, GmCHS produces naringenin chalcone in the absence of GmCHR, which then converts to naringenin, 2-hydroxynaringenin, and genistein by GmCHI and GmIFS (Fig. 1). Multiple binary PPIs have been reported between GmIFS2, which is an active GmIFS isozyme highly associated with (deoxy)isoflavonoid accumulation, and specific isoforms of upstream enzymes, including GmCHS7^17^, GmCHR5^18^, and GmCHI1B1^17^, respectively. Nevertheless, the metabolic outcomes of these PPIs remain unknown. Here, we leveraged yeast as a versatile platform to reconstitute various complex variants using the interacting isoenzymes mentioned above (i.e., GmIFS2, GmCHS7, GmCHR5 and GmCHI1B1) and evaluated the effect of the GmIFS2-mediated PPIs on (deoxy)isoflavonoid biosynthesis. In vivo characterization in yeast via metabolite quantification demonstrated that the inactive GmIFS2 (due to the lack of corresponding CPR in yeast) could enhance GmCHS7 activity via binary PPI in the presence of ER. GmIFS2 could also redirect more metabolic flux toward the deoxyisoflavonoid pathway branch by organizing GmCHS7 and GmCHR5 in close proximity to enhance intermediate diffusion. Conversely, the presence of GmCHI1B1 with inactive GmIFS2 and GmCHS7 augmented the production of naringenin chalcone and naringenin, thereby competing with the deoxyisoflavonoid pathway. We further identified two new isoflavone *O*-methyltransferases (GmIOMTs) that interacted with GmIFS2. The PPIs also regulated the metabolic flux between two competing pathways and favored the deoxyisoflavonoid branch. Overall, our yeast-based characterization work reveals that the GmIFS2-mediated PPIs play significant regulatory roles in soybean (deoxy)isoflavonoid metabolism. This proof-of-concept study demonstrates the feasibility of yeast synthetic biology in elucidating the metabolic role of plant enzyme complexes, particularly the challenging membrane-bound complexes mediated by CYPs.

**Figure 1.**
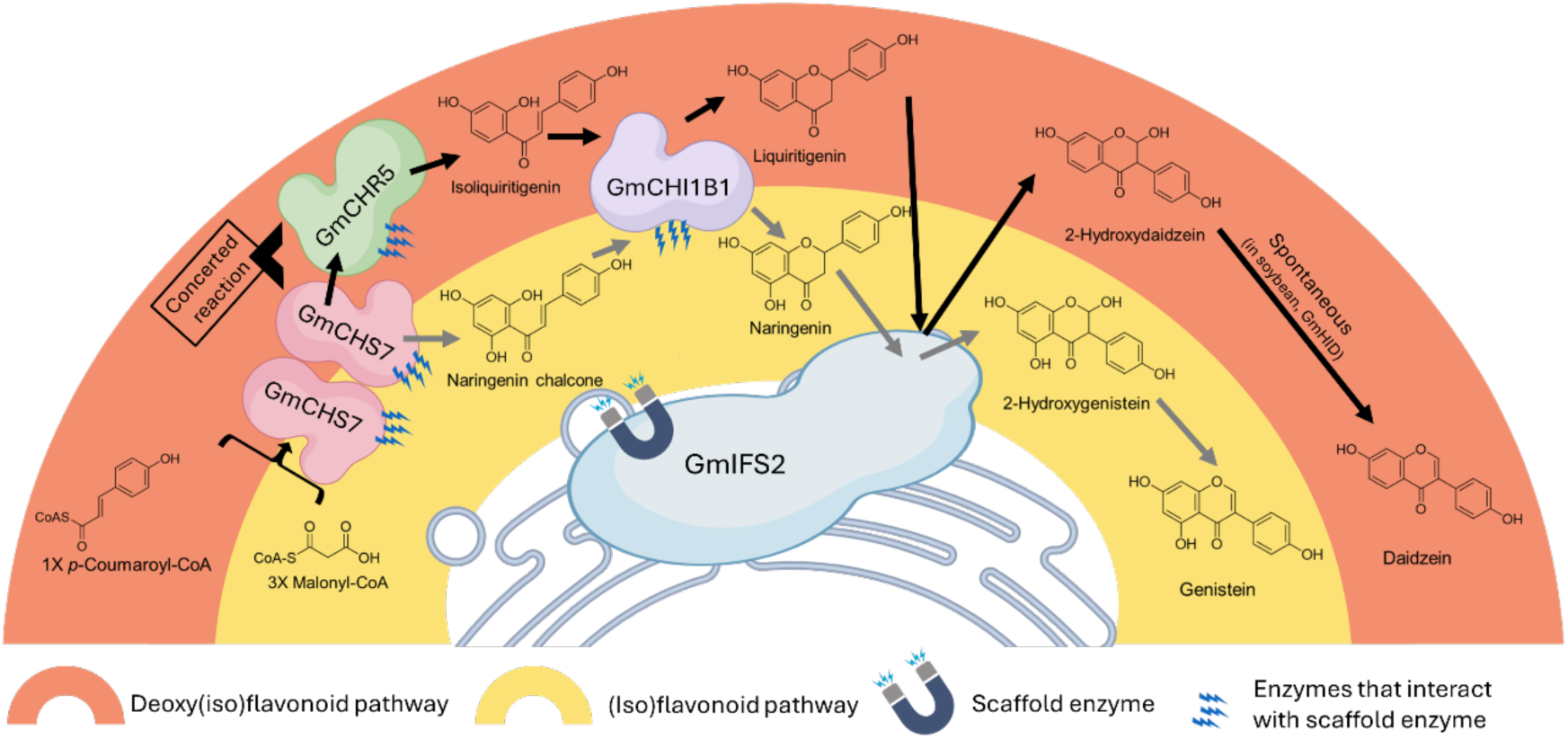
The reconstructed soybean enzyme complex and biosynthetic pathway for the biosynthesis of isoflavonoids, e.g., daidzein and genistein, in yeast. 4CL, 4-Coumarate CoA ligase; CHS, chalcone synthase; CHR, chalcone reductase; CHI, chalcone isomerase; IFS, isoflavone synthase; HID, 2-hydroxyisoflavanone dehydratase. In the enzyme complex, GmIFS2 serves as the scaffold enzyme, while GmCHS7, GmCHR5 and GmCHI1B1 interact with GmIFS2. The biosynthetic pathway branches into two competing pathways, i.e., deoxyisoflavonoid pathway (orange) and isoflavonoid pathway (yellow).

## Results

### Enzyme localization and PPI validation in yeast

The PPIs between the GmIFS2 and the three cytosolic enzymes (i.e., GmCHS7, GmCHR5, and GmCHI1B1) have been validated by yeast 2-hybrid (Y2H) assay^19^ and bimolecular fluorescence complementation (BiFC) in *N. benthamiana* or co-immunoprecipitation in soybean hairy roots^17^ in prior studies. To prove that the soybean enzymes also interact in yeast within similar subcellular localization, we examined the localization of codon-optimized GmIFS2, GmCHS7, GmCHR5, and GmCHI1B1 in yeast using fluorescent fusion proteins. Microscopic analysis has validated that GmIFS2 was targeted to yeast ER, while other enzymes localized in the cytoplasm (Fig. 2A and Supplementary Fig. S1). The PPI between GmIFS2 and each cytosolic enzyme was validated in yeast by BiFC (Fig. 2B) and Y2H (Extended Data Fig. 1). 3’-hydroxy-N-methylcoclaurine 4’-O-methyltransferase of opium poppy (*Papaver somniferum*) (Ps4’OMT)^20^ was used as the negative control in the PPI assays (Supplementary Fig. S2). Furthermore, the co-localization of GmIFS2-eCFP and GmCHS7-, GmCHR5-, or GmCHI1B1-eGFP proved that the PPI took place on the ER membrane, with GmHID downstream of GmIFS2 used as the negative control (Fig. 2C). There were no obvious signs of binary PPIs among GmCHS7, GmCHR5, and GmCHI1B1, consistent with previous reports (Extended Data Fig. 2).

**Figure 2.**
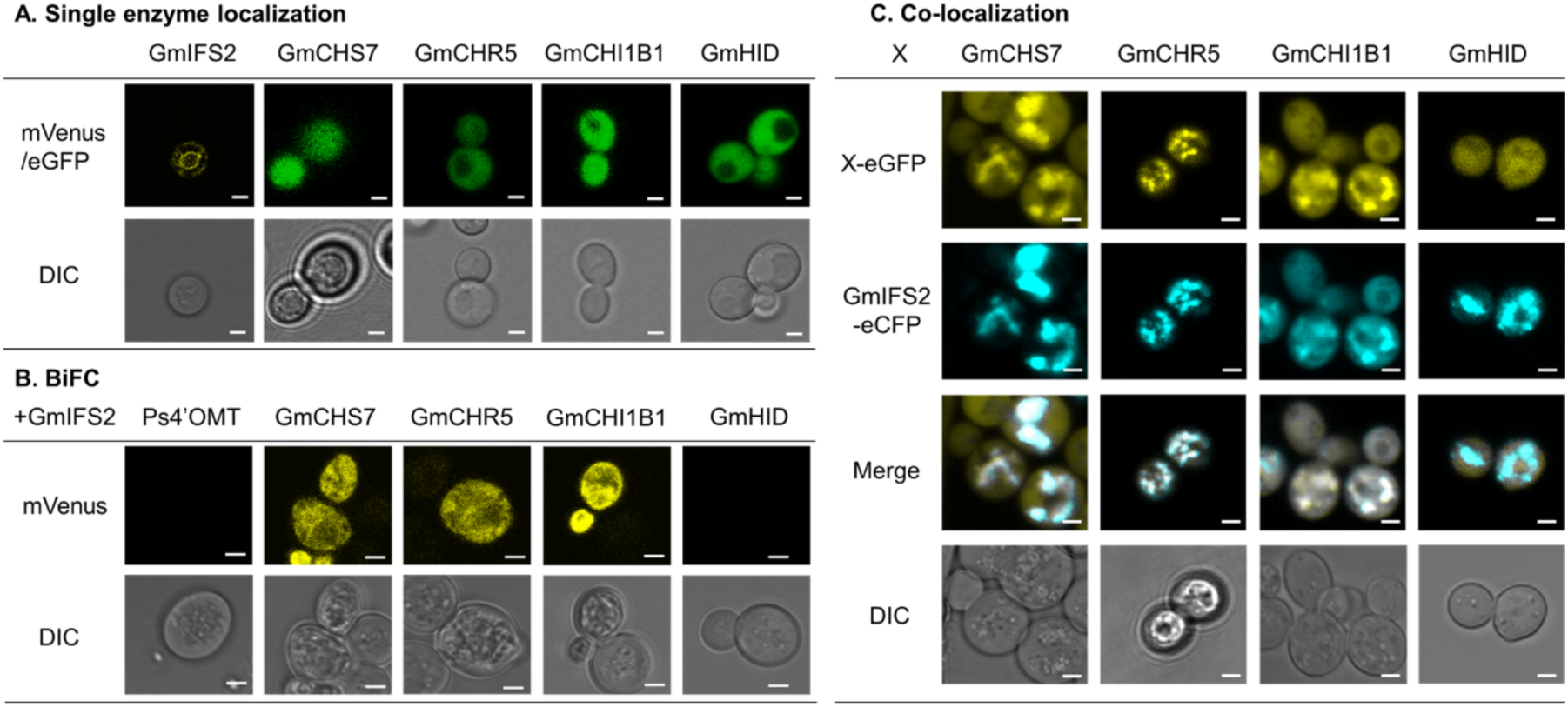
Soybean isoflavone enzyme localization and PPI validation in yeast. **(A)** Confocal microscopic analysis of single enzyme localization in yeast. GmIFS2 was fused with mVenus at the C-terminus. GmCHS7, GmCHR5, GmCHI1B1 and GmHID were fused with enhanced green fluorescent protein (eGFP) at the C-terminus. **(B)** Detection of binary interactions of GmIFS2 with GmCHS7, GmCHI1B1, GmCHR5 and GmHID by BiFC in yeast cells. mVenus protein was split into C-terminal (CV) and N-terminal (NV) parts. GmIFS2-CV and X-NV or NV-X were co-expressed in yeast cells, where X refers to GmCHS7, GmCHR5, GmCHI1B1, GmHID and Ps4’OMT. **(C)** Co-localization assay of GmIFS2 with GmCHS7, GmCHI1B1, GmCHR5 and GmHID in yeast cells. GmIFS2 was fused with enhanced cyan fluorescent protein (eCFP) at the C-terminus. GmCHS7, GmCHB1, GmCHR5 and GmHID (negative control) were fused with eGFP at the C-terminus. False colors were used for mVenus, eCFP, and eGFP. Cell morphology is observed with differential interference contrast (DIC). Scale bars = 2 μm.

### Evaluating the metabolic impact of GmIFS2-mediated binary PPIs in yeast revealed enhanced productivity of GmCHS7

We developed a yeast platform to evaluate the metabolic effects of the binary PPIs between GmIFS2 and its interacting cytosolic enzymes. Evaluating this effect in the native host or in *N. benthamiana* would be challenging, because a functional GmIFS2 would unavoidably accelerate the conversion of interacting enzymes’ products directly (e.g., liquiritigenin) or indirectly (e.g., isoliquiritigenin) to downstream 2-hydroxyisoflavanones. Therefore, we expressed GmIFS2 and its interacting enzymes in a yeast CEN.PK2-1D^8^ without plant CYPR to inhibit the biochemical reaction catalyzed by GmIFS2 (Extended Data Fig. 3).

Firstly, we integrated both GmIFS2 and GmCHS7 into the yeast genome to evaluate the effect of PPI on GmCHS7 activity. A gene encoding 4-coumarate: CoA ligase (Gm4CL3) was also integrated to produce 4-coumaroyl-CoA from fed *p*-coumaric acid in vivo, yielding an engineered strain named yCL1 that produces naringenin chalcone from coumaroyl-CoA and endogenous malonyl-CoA. We also developed a control strain (yCL2) without PPI by replacing GmIFS2 with the fluorescent protein mCherry. Both strains were cultured in synthetic dropout (SD) medium overnight to saturation, and then supplemented with 2 mM *p*-coumaric acid for 48 hours. As naringenin chalcone can convert to naringenin spontaneously in yeast and in planta^21^, both naringenin chalcone and naringenin in the supernatant were quantified by liquid chromatography mass spectrometry (LC-MS) to evaluate the activity of GmCHS7. We found that the presence of GmIFS2 enhanced the production of naringenin chalcone (referred to as NC) and naringenin (N) by 41.6% from 57.18 μM in yCL2 to 80.98 μM in yCL1 (six independent replicates, P<0.0001, Fig. 3A). Only trace amount of GmIFS2 downstream products, including 2-hydroxygenistein and genistein were detected (0.96 μM, less than 1.2% of the total titer of NC and N), proving that the endogenous CPR in yeast is insufficient for the functioning of GmIFS2. We then tuned the expression level of GmIFS2 in yeast by expressing GmIFS2 in low-copy and high-copy plasmids^22^, respectively, with GmCHS7 integrated into the yeast genome, and used mCherry to replace GmIFS2 as the negative controls. We found the production enhancement correlated with the expression level of GmIFS2. Higher expression of GmIFS2 increased GmCHS7 products by 69.0%, while lower expression of GmIFS2 increased by 34.2% (Fig. 3B). To exclude the possible ER disturbance-related impacts after expressing any CYPs, we replaced the GmIFS2 in yCL1 with an unrelated plant CYP, canadine synthase (CjCAS) from *Coptis Japonica*^23^. The production of NC+N in this control strain showed no significant difference to yCL2 (Extended Data Fig. 4). Taken together, GmIFS2 could increase the catalytic activity of GmCHS7 solely by PPI, with a higher GmIFS2 expression level leading to greater enhancement. The involvement of the ER was also important to the production enhancement. When GmIFS2 was re-localized to cytoplasm (Extended Data Fig. 5) by truncated 25 residues at N-terminal (sequence shown in Supplementary Table S1), the production of naringenin chalcone and naringenin dropped to 29.19 μM, lower than that in both yCL2 and yCL1, despite that the cytosolic GmIFS2 still interact with GmCHS7 (Extended Data Fig. 6). This decrease highlighted the importance of the ER in regulation mediated by complexes.

**Figure 3.**
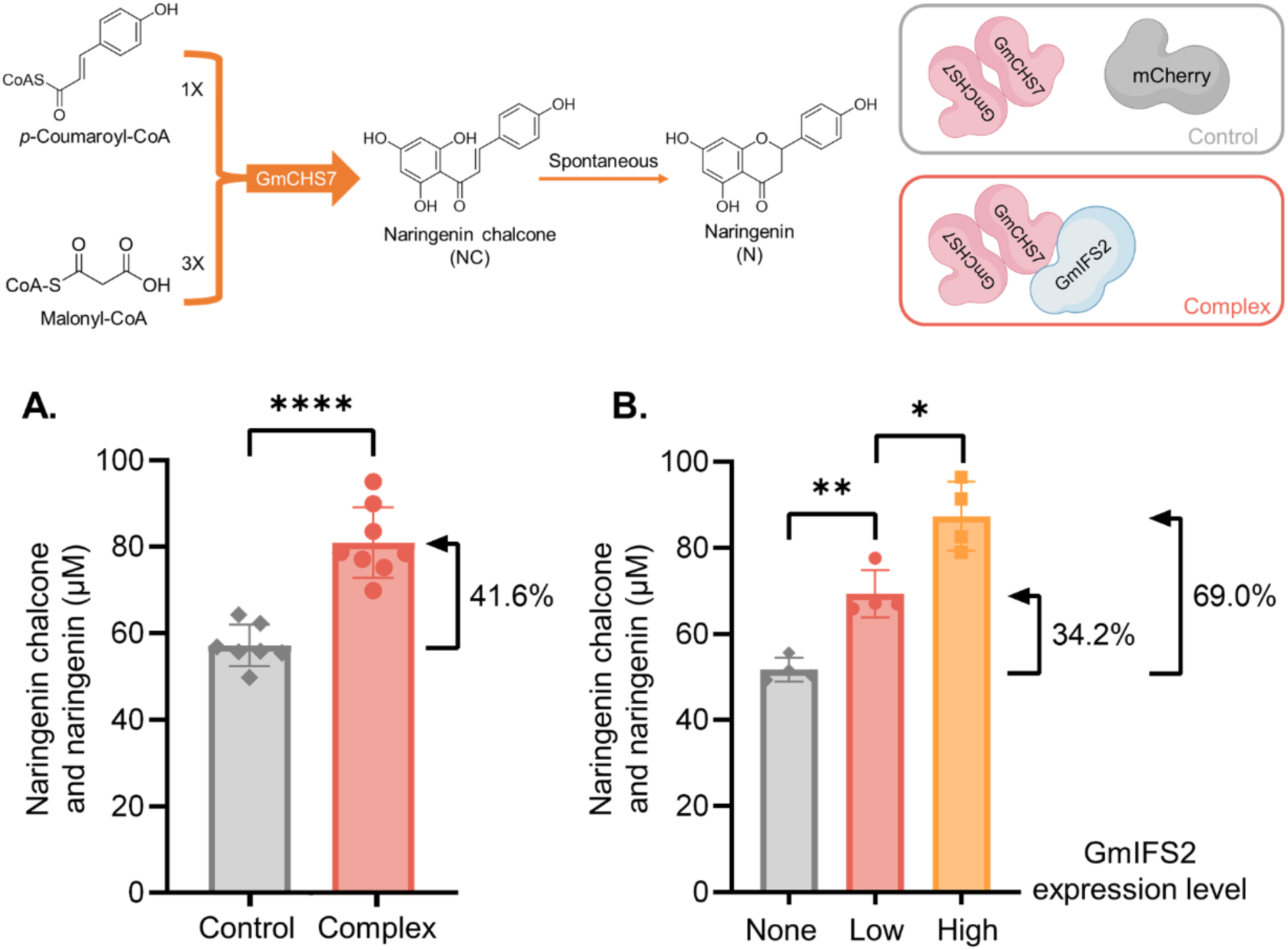
The biochemical effects of GmIFS2 on the activity of GmCHS7 via PPI in yeast. **(A)** Production of naringenin chalcone and naringenin by yeast strains harboring enzyme complexes (inactive GmIFS2 with GmCHS7) or control (mCherry with GmCHS7). Data are means ± SD for six independent clones (t test, ****P < 0.0001). **(B)** Production of naringenin chalcone and naringenin by yeast strains with different GmIFS2 expression levels. Data are means ± SD for four independent clones (t test, *P < 0.05, **P < 0.01).

We also investigated the possible metabolic effect of the PPI between GmIFS2 and GmCHI1B1 by co-expressing inactive GmIFS2 or mCherry with GmCHI1B1 and supplementing 80 μM of naringenin chalcone or isoliquiritigenin to the cultural medium, respectively. After 4 or 24 hours, the titers of corresponding products of GmCHI1B1, including naringenin or liquiritigenin, were quantified by LC-MS and compared (Extended Data Fig. 7). Despite the validated PPI in our and prior studies, the productivity of GmCHI1B1 did not change. We did not evaluate the effect of GmIFS2-mediated PPI on GmCHR5, because its substrate is unstable and can only be synthesized via the concerted reaction catalyzed by GmCHS. Instead, a putative ternary complex containing GmIFS2, GmCHS7, and GmCHR5 was constructed and characterized as described below.

### GmCHI1B1 further augmented the production enhancement of the GmIFS2-GmCHS7 complex

We then investigated the metabolic effect of the PPI between inactive GmIFS2 and GmCHI1B1 in the presence of other cytosolic enzymes. We co-expressed GmCHS7, GmCHI1B1, and inactive GmIFS2 to develop yCL3, which produced 83.68 μM NC+N (eight independent replicates, P<0.0001, Fig. 4A). The titer was 77.5% higher than that of the control strain yCL4 (47.14 μM) that did not contain GmIFS2. As the binary GmIFS2-GmCHS7 complex only enhanced GmCHS7’s biosynthetic activity by 41.6%, the higher enhancement in yCL3 indicated the formation of a ternary GmIFS2-GmCHS7-GmCHI1B1 complex, which further increased GmCHS7’s productivity. We developed an engineered yeast strain co-expressing GmIFS2-DsRed, GmCHS7-eCFP and GmCHI1B1-eGFP and performed time-lapsed microscopic analysis. Results showed that the assembly process took place sequentially in a consecutive manner (Fig. 4B and Video S1). The interaction between GmIFS2 and GmCHS7 took place first, followed by the recruitment of GmCHI1B1 to the ER membrane. The binary PPI between GmIFS2 and GmCHS7 likely forms the core of the GmIFS2-mediated complexes, which is reasonable given that GmCHS7 is crucial for both the flavonoid and deoxyflavonoid pathways and is likely to be tightly regulated.

**Figure 4.**
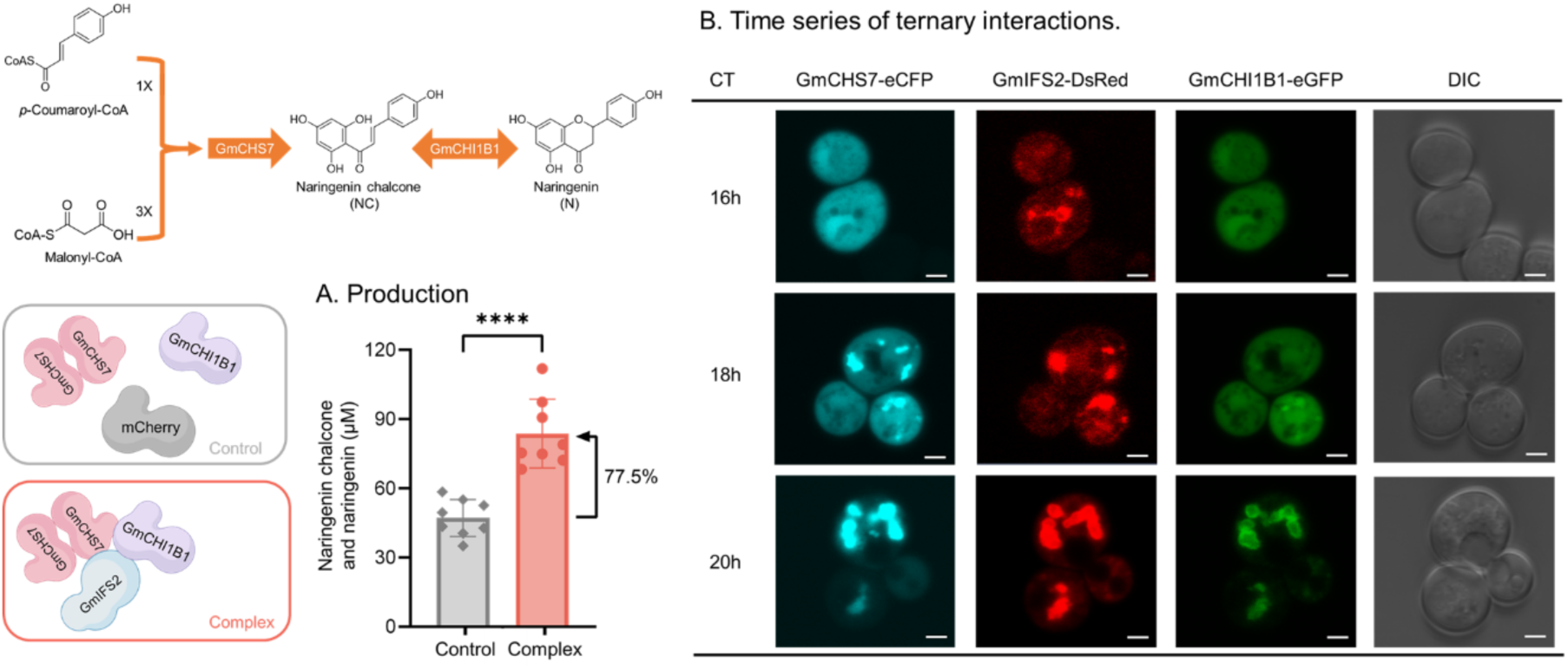
Characterization of the ternary GmIFS2-GmCHS7-GmCHI1B1 complex in yeast. **(A)** Production of naringenin chalcone and naringenin by yeast strains harboring enzyme complexes (inactive GmIFS2 with GmCHS7 and GmCHI1B1) or control (mCherry with GmCHS7 and GmCHI1B1). 2 mM of p-coumaric acid was fed and the supernatant was analyzed after 48 hours. Data are means ± SD for eight independent clones (t test, ****P < 0.0001). **(B)** Ternary co-localization observed under microscopy. The cells were co-expressed with GmIFS2-DsRed, GmCHS7-eCFP and GmCHI1B1-eGFP. Observation was performed at cell culture time (CT) 16 h, 18 h and 20 h respectively. Intracellular fluorescence within a single focal plane in the yeast cells was observed by means of confocal laser fluorescent microscopy. The excitation wavelength of eCFP was 405 nm, and the detection wavelength of eCFP was 447-498 nm. The excitation wavelength of eGFP was 488 nm, and the detection wavelength of eGFP was 500-549 nm. The excitation wavelength of DsRed was 561 nm, and the detection wavelength of DsRed was 564-703 nm. eCFP used the false color cyan; eGFP used the false color green; DsRed used the false color red. Cell morphology is observed with differential interference contrast (DIC). Scale bars = 2 μm.

### Assembly of the possible ternary GmIFS2-GmCHS7-GmCHR5 complex redirects more metabolic flux toward deoxyflavonoid production

We evaluated the effect of GmIFS2-mediated PPIs in the presence of both GmCHS7 and GmCHR5, focusing on their possible role on tuning the metabolic flux between the two competing pathway branches producing deoxyflavonoid and flavonoid. In the deoxyflavonoid branch, GmCHR5 accepts the unstable coumaryl-trione produced by GmCHS7 and converts it to isoliquiritigenin^12^. The deoxyflavonoid pathway branch involves two concerted reactions catalyzed by GmCHS7 and GmCHR5 and competes with the flavonoid branch, in which coumaryl-trione aromatizes to naringenin chalcone. It has long been postulated that GmCHS7 and GmCHR5 interact directly or indirectly to facilitate intermediate diffusion.

In this test, we co-expressed GmCHS7 with GmCHR5 to provide the appropriate substrate in vivo. Additional integration of GmIFS2 or mCherry led to engineered strains yCL5 and yCL6, respectively. Both strains contained two competing pathway branches, including the GmCHS7-catalyzed branch toward naringenin chalcone (NC) and the branch comprised of both GmCHS7 and GmCHR5 toward isoliquritigenin (IL). We evaluated the product preference of these pathways based on the ratio of IL and NC. Their spontaneous downstream products, namely naringenin (N) and liquiritigenin (L), were also considered. In the absence of the complex, yCL6 produced 9.9 μM flavonoids (NC+N) and 33.2 μM deoxyflavonoids (IL+L) (Fig. 5A), leading to the product ratio (R = (IL+L) : (NC+C)) of 3.35 (Fig. 5B). The presence of GmIFS2 and consequent formation of the ternary GmIFS2-GmCHS7-GmCHR5 complex in yCL5 increased the ratio to 4.32 (Fig. 5B). The titers of NC+C and IL+L in yCL5 were 14.9 μM (50.5% increase) and 64.3 μM (93.6% increase), respectively (Fig. 5A). While the enhancement of the overall titer was likely due to the PPI between GmIFS2 and GmCHS7, the change in the IL+L and NC+C ratio from 3.35 to 4.32 indicated that the formation of the ternary complex redirected more metabolic flux toward the deoxyflavonoid pathway branch. We also analyzed the yCL5 and yCL6 strains during a 72-hour fermentation and observed the enhancement as early as 24 hours after cultivation began, indicating that the complex assembly was completed by then (Extended Data Fig. 8).

**Figure 5.**
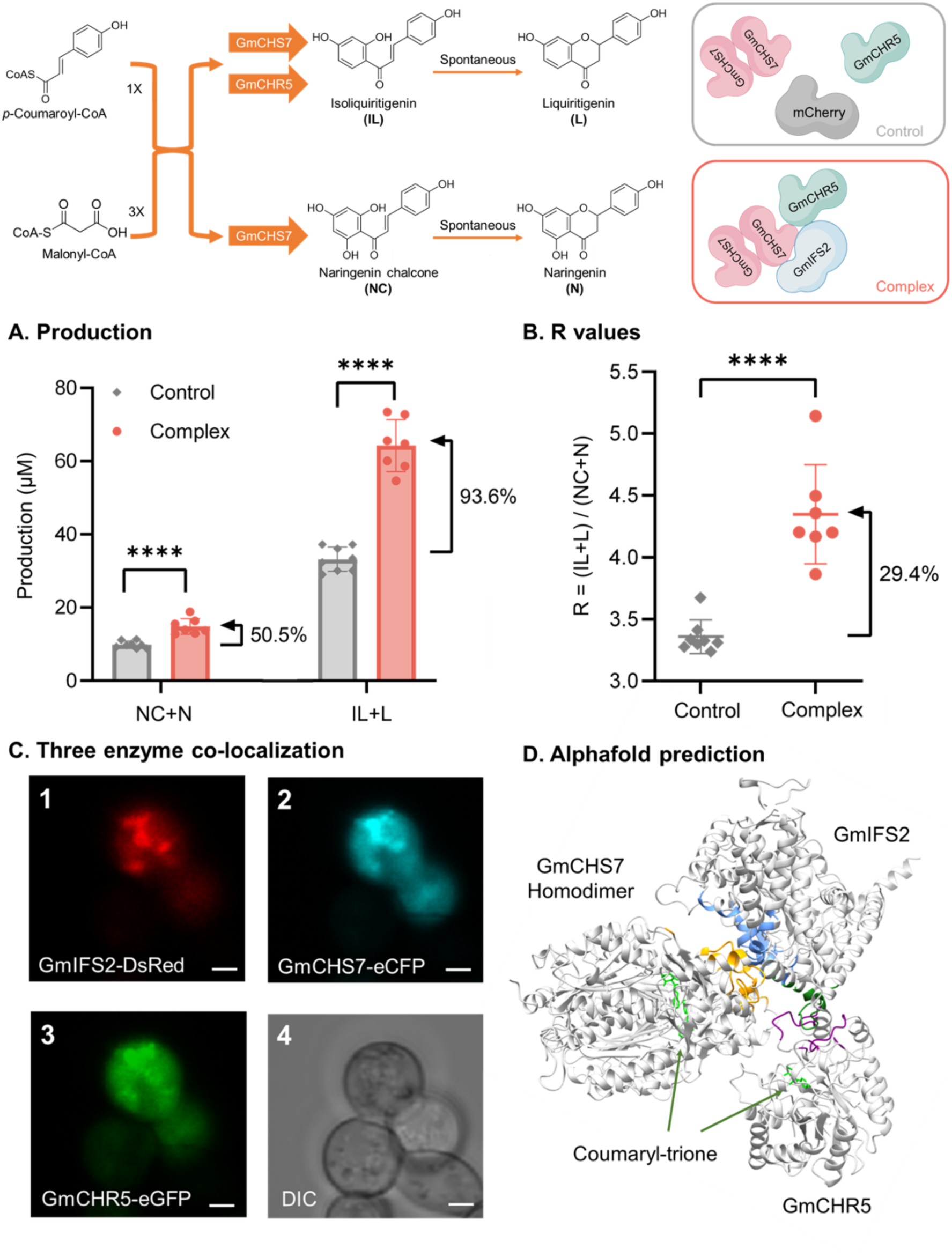
Characterization of the ternary GmIFS2-GmCHS7-GmCHR5 complex in yeast. **(A)** Production of naringenin chalcone and naringenin (NC+N) and isoliquiritigenin and liquiritigenin (IL+L) by yeast strains harboring enzyme complexes (inactive GmIFS2 with GmCHS7 and GmCHR5) or control (mCherry with GmCHS7 and GmCHR5). Data are means ± SD for six independent clones (t test, ****P < 0.0001). **(B)** R values of control strain and complex strain by calculating IL+L production over NC+N production. **(C)** Three enzyme co-localization detection by co-expressing their fluorescence-protein chimeras in yeast cells. GmIFS2, GmCHS7 and GmCHR5 were fused with DsRed, eCFP and eGFP at C-terminus, respectively. Intracellular fluorescence within a single focal plane in the yeast cells was observed by means of confocal laser fluorescent microscopy. Scale bars = 2 μm. **(D)** The ternary enzyme complex structure predicted by Alphafold-Multimer. Amino residues within 5 Å to other proteins were identified as the interface of the protein complex. Interacting residues in GmIFS2, GmCHS7 and GmCHR5 were labeled in blue, yellow and purple, respectively. Docking of substrate p-coumarylcyclohexantrion (as shown in green) in GmCHS7 and GmCHR5 was performed in Autodock.

There was no evidence indicating the direct PPIs between GmCHS7 and GmCHR5, regardless of the presence of GmIFS2 (Extended Data Fig. 2 and Fig. 5D). Instead, the enhanced production of isoliquiritigenin is likely due to GmIFS2 assembling GmCHS7 and GmCHR5 and recruiting them in closer proximity. We proposed that either GmIFS2, GmCHS7 and GmCHR5 form a ternary complex, or they interact via binary PPIs (GmIFS2-GmCHS7 and GmIFS2-GmCHR5) but both on ER membrane. Both hypothetical mechanisms can lead to the proximity of GmCHS7 and GmCHR5. We performed a ternary co-localization assay of GmIFS2-DsRed, GmCHS7-eCFP and GmCHR5-eGFP, showing that GmCHS7 and GmCHR5 can be recruited on the ER membrane simultaneously in the presence of GmIFS2 (Fig. 5C). An AlphaFold-Multimer model^24^ of the ternary complex supported the first hypothesis of a ternary complex, in which GmCHS7 and GmCHR5 interact with two distinct but spatially close regions on GmIFS2 (Lys90-Ser100/Asn141-Arg152 for GmCHS7 and Asp282-Thr290 for GmCHR5) (Fig. 5D).

### Biochemical characterization suggests competition between GmCHI1B1 and GmCHR5

To investigate the possible metabolic role and synergistic effect of a putative quaternary complex, we developed yCL7 containing inactive GmIFS2, GmCHS7, GmCHR5, and GmCHI1B1. However, the engineered strain did not further favor deoxyflavonoid production or enhance the total titer of all products derived from GmCHS7, suggesting that GmCHI1B1 and GmCHR5 might not interact with GmIFS2 at the same time. The ratio of IL+L and NC+N dropped from 3.86 in the control strain yCL8 to 2.83 in yCL7 (26.8% decrease) (Fig. 6B), opposite to the enhancement from 3.35 to 4.32 in yCL5 containing GmIFS2-GmCHS7-GmCHR5(Fig. 5B).

**Figure 6.**
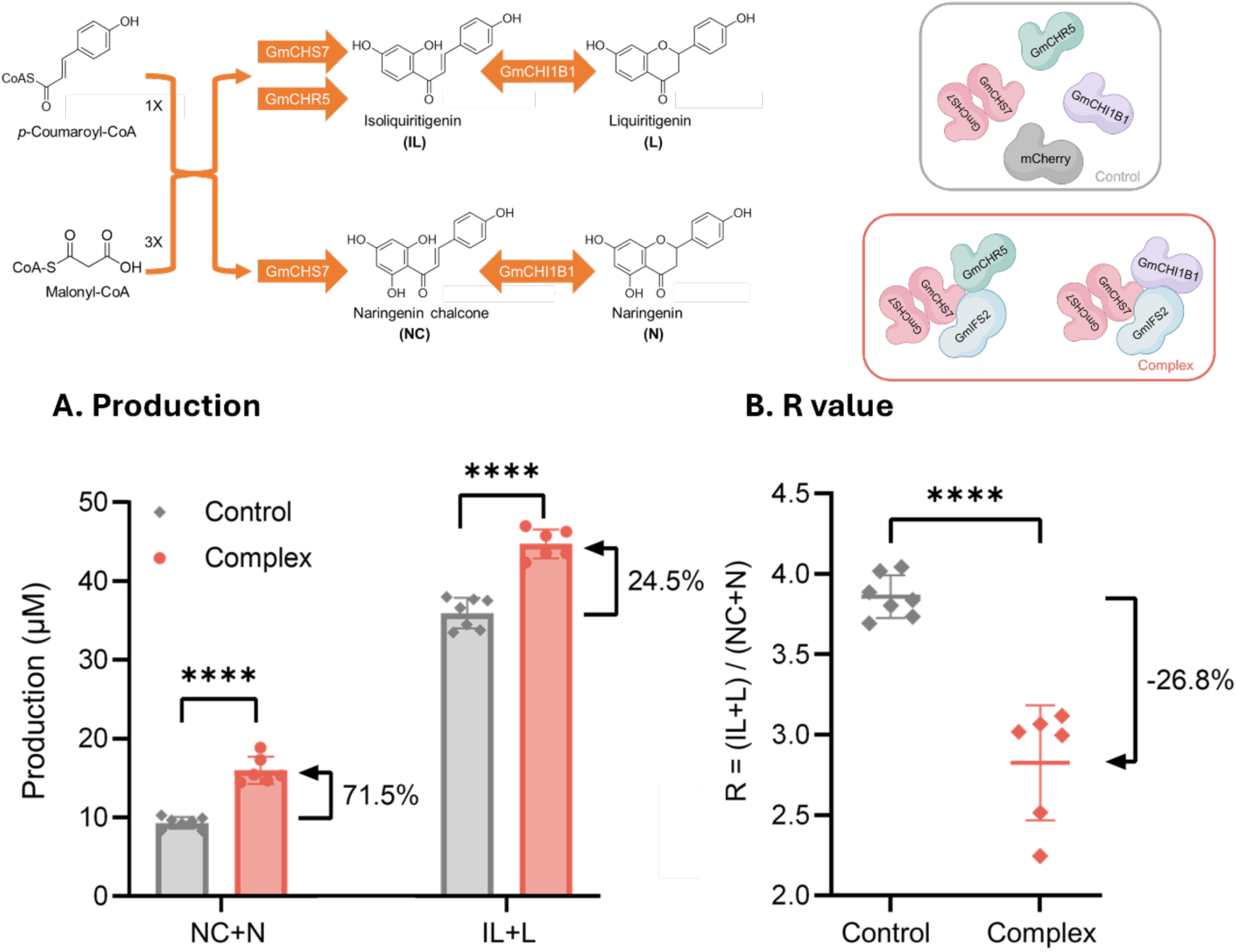
Coexpressing inactive GmIFS2, GmCHS7, GmCHI1B1 and GmCHR5 in yeast. **(A)** Production of naringenin chalcone and naringenin (NC+N) and isoliquiritigenin and liquiritigenin (IL+L) by yeast strains harboring enzyme complexes (inactive GmIFS2 with GmCHS7, GmCHR5 and GmCHI1B1) or control (mCherry with GmCHS7, GmCHR5 and GmCHI1B1). Data are means± SD for six independent clones (t test, ****P < 0.0001). **(B)** R values of control strain and complex strain by calculating IL+L production over NC+N production.

The decreased ratio resulted from the inconsistent titer enhancement in yCL7. While the titer of NC+C increased by 71.5% from 9.33 μM in yCL8 to 16.00 μM in yCL7, the titer of IL+L only increased by 24.5% from 35.94 μM in yCL8 to 44.73 μM in yCL7 (six independent replicates, P<0.0001, Fig. 6A). It is plausible that GmCHI1B1 blocks the PPI between GmCHR5 and GmIFS2 by occupying the same space and causing steric hindrance. An AlphaFold-Multimer model^24^ supported this hypothesis, showing that GmCHI1B1 and GmCHR5 likely compete when interacting with GmIFS2 (Extended Data Fig. 9). Consequently, there might be two competing ternary complexes co-existing in yCL7, including the GmIFS2-GmCHS7-GmCHR5 complex that favors the deoxyflavonoid pathway branch and the GmIFSs-GmCHS7-GmCHI1B1 complex that solely enhances flavonoid pathway branch. Notably, the overall titers from yCL5 to yCL8 did not surpass those in yCL1 or yCL2, likely due to the higher expression burden or the limited supply of upstream precursors.

### GmIFS2 modulates the substrate preference of GmIOMTs toward the deoxyflavonoid branch via PPI

Given the distinct product preferences of GmIFS2-mediated ternary complexes, we hypothesized that GmIFS2-mediated PPIs are extensive in (deoxy)isoflavonoid metabolism and used the established yeast platform to explore their metabolic effects. We selected isoflavone *O*-methyltransferases, which catalyze daidzein and genistein to formononetin and biochanin A, respectively. Previous research only demonstrated indirect association between IFS and IOMT in alfalfa^25^, and only one soybean IOMT (encoded by *Glyma05g14700*^26^) was characterized with no validated PPIs. We firstly searched for more putative soybean IOMTs via gene mining^27^ and used yeast for biochemical characterization and subsequent PPI identification. We identified and cloned eight putative IOMT genes from soybean. The putative genes (i.e., *Glyma08g27260, Glyma10g32030, Glyma13g24210, Glyma18g50260, Glyma18g50280, Glyma18g50290, Glyma20g35610,* and *Glyma20g35630*) were renamed as *GmIOMT8, IOMT10, IOMT13, IOMT18-1, IOMT18-2, IOMT18-3, IOMT20-1,* and *IOMT20-2,* respectively (Supplementary Table S1). Biochemical characterization in yeast via GmIOMT expression and substrate feeding identified six functional GmIOMTs with varied substrate preference. GmIOMT8, 18-1, 18-2, 18-3, and 20-1 can *O*-methylate the 4’-site of isoflavones, producing formononetin from fed daidzein or biochanin A from fed genistein (Figure 7A). GmIOMT13 exhibited a different substrate preference, methylating the 4’-hydroxyl group of naringenin or liquiritigenin toward methylated naringenin (m/z 271.0965, 20ppm) or methylated liquiritigenin (m/z 287.0914, 20ppm), respectively (Figure 7A). GmIOMT10 and GmIOMT20-2 did not show detectable methylation activity on naringenin, liquiritigenin, genistein, or daidzein.

**Figure 7.**
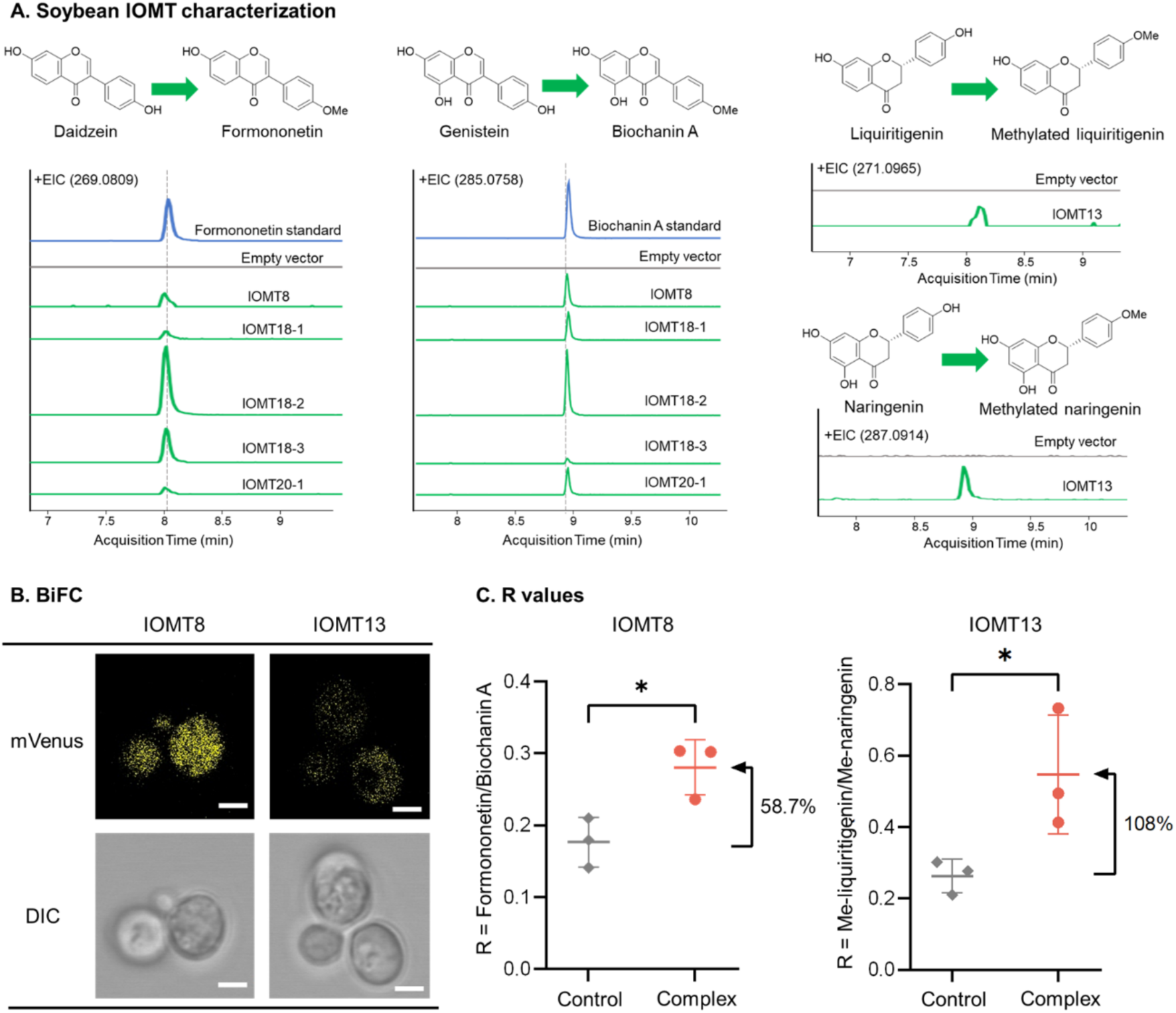
Characterization of GmIOMTs’ activities and PPIs with GmIFS2. **(A)** Functional characterization of GmIOMTs in yeast. Extracted ion chromatogram (EIC) of methylated products are shown. Traces are representative of three biological replicates. **(B)** Detection of binary interactions of GmIOMTs with GmIFS2 by BiFC in yeast cells. BiFC test showed the interaction of GmIFS2 with GmIOMT8 and GmIFS2 with GmIOMT13. **(C)** R ratios of control strain and complex strain by calculating formononetin production over biochanin A production, or methylated liquiritigenin production over methylated naringenin production. Data are means ± SD for at three independent clones (t test, *P < 0.05).

We then screened the possible PPIs between the GmIOMTs with GmIFS2 by yeast-based BiFC and Y2H and identified two interacting enzymes, including GmIOMT8 and GmIOMT13 (Fig. 7B and Extended Data Fig. 10). The metabolic effect of these PPIs was then evaluated in yeast. Yeast harboring GmIOMTs, with or without GmIFS2, were cultured in SD-ura-trp medium supplemented with 50 μM naringenin, liquiritigenin, genistein, or daidzein, respectively. We use the ratio of peak area of deoxyisoflavonoid and isoflavonoid products in LC-MS to evaluate their preference of the two branches, given the lack of a full set of chemical standards for all products. Co-expressing GmIFS2 with GmIOMT8 or GmIOMT13 redirects more flux toward the deoxyisoflavonoid pathway branches (Fig. 7C) by specifically inhibiting the isoflavonoid branches (Extended Data Fig. 10). In contrast, the preference of the other four non-interacting GmIOMTs did not change (Extended Data Fig. 10). The ratio of formononetin and biochanin A produced by GmIOMT8 increased from 0.18 to 0.28 (Fig. 7C). Similarly, when GmIFS2 was co-expressed with GmIOMT13, the ratio of methylated liquiritigenin and methylated naringenin increased from 0.26 to 0.54 (Fig. 7C). Protein structural simulation and substrate docking results suggested a loop of GmIFS2 (from Phe217 to Gly227) inserts into the interface of the GmIOMT8 dimer. Since the interface forms the back wall of the substrate binding site, the insertion of GmIFS2 loop can restrict the binding pocket, thus altering GmIOMT’s substrate preference (Extended Data Fig. 10). These results indicate that GmIFS2 can direct metabolic flux to the deoxyisoflavonoid branch by altering the original substrate preferences of GmIOMT8 and GmIOMT13 via PPI.

## Discussion

In this study, we developed a yeast platform to elucidate the regulatory roles of CYP-mediated plant enzyme complexes in vivo, demonstrated by characterizing the metabolic effects of GmIFS2-mediated PPIs in (deoxy)isoflavonoid biosynthesis. To decouple the metabolic effect of the binary PPIs from the catalytic functions from the two interacting enzymes, we used an inactive GmIFS2 to evaluate how GmIFS2 altered the metabolic products of cytosolic enzymes (e.g., GmCHS7, GmCHR5, and GmCHI1B1) and expanded the evaluation to two newly identified GmIOMTs.

Our study highlighted the feasibility of the yeast platform to reconstruct and characterize CYP-centered plant enzyme complexes. The shorter growth time and higher efficiency of yeast in PPI identification and biochemical characterization enable rapid elucidation of the metabolic functions of complexes. Particularly, the membrane-bound organelles such as ER in yeast allows for in vivo characterization of these complexes, providing an alternative approach to the traditional method that isolates enzyme complexes directly from native plants. Although isolation methods have proven effective for harvesting the CYP-mediated dhurrin complex from sorghum lipid particles^28^, many membrane-bound enzyme complexes are fragile and cannot not be isolated properly. Notably, previous co-immunoprecipitation efforts using soybean could not recover GmCHS7 or GmCHR5 interacting with GmIFS2^17^ despite other validation of these PPIs, highlighting the need for a yeast platform for complex characterization. The lack of functional CPRs and the simpler, orthogonal metabolic context in yeast allow for evaluating the effect of CYP-mediated PPIs on PNP production. The insufficient CPR supply decouples the effect of CYP-mediated PPI from CYP’s own catalytic activity, enabling subsequent functional characterization. Limited crosstalk between plant CYPs and yeast endogenous metabolism is also beneficial for complex characterization. In contrast, when we reconstructed the GmIFS2-mediated complexes in the model plant *Nicotiana benthamiana*, we did not observe any (deoxy)isoflavonoid products (data not shown). Similar results were observed in *N. tabacum* leaf transformed with GmIFS1^29^, possibly due to crosstalk between the heterologous soybean pathway and the native metabolism in *Nicotiana* plants.

The metabolic roles of PPIs elucidated in yeast provide new insights into the plant’s post-translational regulation mechanism. Unlike the large enzyme complexes identified in primary metabolism that involves a complicated PPI network (e.g., the metabolon in the tricarboxylic acid cycle^30,31^), the PPIs regulating (deoxy)isoflavonoid synthesis are all mediated by GmIFS2 and are likely binary. The metabolic effect of the binary PPI was demonstrated by the enhancement of GmCHS7’s biosynthetic activity, likely due to the re-localization of GmCHS7 from cytoplasm to ER. It has been reported that malonyl-CoA is naturally enriched around the ER for optimal lipid biosynthesis and fatty acid elongation^32,33^. Additionally, the lipid-rich ER membrane can facilitate the movement of hydrophobic intermediates along the ER’s outer face to increase local malonyl-CoA concentration. A similar mechanism has been proposed in the C4H-mediated complex to enable cinnamic acid channeling for enhanced phenylpropanoid biosynthesis^34^. The enhancement of GmCHS7’s productivity might result from the same mechanism. Re-localizing the binary GmIFS2-GmCHS7 complex to the cytoplasm significantly lowered the production of naringenin chalcone, further supporting this hypothesis (Extended Data Fig. 3).

Given the limited surface space of GmIFS2 for interaction and the possible steric hindrance, the number of cytosolic enzymes that GmIFS2 can assemble simultaneously is likely limited. Our characterization results indicate the existence of two competing ternary enzyme complexes in yeast, in which GmIFS2 can interact with GmCHS7 and GmCHR5, or GmCHS7 and GmCHI1B1, respectively. The core GmIFS2-GmCHS7 interaction persists in both complex and continues augmenting GmCHS7’s productivity in the presence of GmCHR5 or GmCHI1B1. Importantly, each of the two competing complexes favors a different, competing pathway branch, respectively. The GmIFS2-GmCHS7-GmCHR5 complex favors deoxyflavonoid (i.e., isoliquiritigenin and downstream liquiritigenin) production by facilitating the diffusion of more pathway intermediates to GmCHR5, which is likely important to the biosynthesis of downstream deoxyisoflavonoids in soybean. As GmCHR5 only acts on the unstable intermediate coumaryl-trione produced by GmCHS7 during the multi-step reaction toward naringenin chalcone^12,18^, the production of isoliquiritigenin catalyzed by GmCHS7 and GmCHR5 competes with the GmCHS7-catalyzed flavonoid pathway unavoidably. It has long been postulated that legume CHS and CHR must catalyze isoliquiritigenin synthesis in a concerted way^12,18^. However, both structural analysis^35^ and PPI identification efforts^18^ have proven that direct PPI between legume CHS and CHR is highly unlikely. In vitro assay has indicated indirect PPI when both GmCHS7 and GmCHR5 were co-immobilized to Ni^2+^-coated magnetic beads and enhanced isoliquiritigenin production^18^. Here, our yeast-based assay provided the first in vivo evidence showing that the interaction between GmCHS7 and GmCHR5 is facilitated by individual assembly on the GmIFS2 scaffold. The increased proximity between GmCHS7 and GmCHR5 might facilitate the transfer of coumaryl-trione from GmCHS7 to GmCHR5 toward isoliquiritigenin biosynthesis, which otherwise would be converted to naringenin chalcone directly. Meanwhile, the GmIFS2-GmCHS7-GmCHI1B1 complex favors flavonoid (i.e., naringenin chalcone and naringenin) production by both augmenting the metabolic flux toward naringenin chalcone and blocking the assembly of GmCHR5. Both biochemical characterization and interaction models suggest that the formation of a bigger quaternary enzyme complex is unlikely in yeast or plants due to the possible competition between GmCHI1B1 and GmCHR5.

Prior research on the regulatory mechanism in soybean (deoxy)isoflavonoid metabolism has been focused on the spatiotemporal differential expression of isozymes. For example, GmIFS2 is highly expressed in root, the primary site for (deoxy)isoflavonoid synthesis^36^, and is induced by stress^18,37–40^, while its isozyme GmIFS1 is ubiquitously expressed^41^. GmCHS7 is also mainly expressed in root^37,42^. Additionally, the expression level of GmCHR5 is at least 10-fold higher than other functional isozymes including GmCHR1 and GmCHR6^18^. GmCHI1B1 is also highly expressed in root^40^. The transcriptional regulation highly correlates with (deoxy)isoflavonoid production. However, the regulatory machinery to tune the metabolic flux between the deoxyisoflavonoid pathway branch and the isoflavonoid branch was not fully identified. Particularly, only the deoxyisoflavonoid pathway can lead to the production of glyceollins^43^, which are an important family of defensive chemicals produced by soybean in response to microbial infections. It is plausible that the GmIFS2-mediated complexes assemble transiently at different time points and under different stresses in planta, thereby dynamically redirecting more metabolic flux toward isoliquiritigenin and downstream glyceollin production when necessary.

Notably, there are still open questions yet to be addressed. Although characterizing PPIs’ metabolic functions in vitro or in a heterologous host has proven effective in revealing their functions in the native host^2^, the ratio of interacting enzymes might vary^2,44^. We have shown that increasing the expression level of GmIFS2 in yeast can further enhance GmCHS7’s biosynthetic activity, but the ratio of GmIFS2 and GmCHS7 in different soybean tissues and the corresponding enhancement effect remain unknown. Moreover, the discovery of new GmIOMTs interacting with GmIFS2 has proven the existence of other complexes in soybean (deoxy)isoflavonoid metabolism, but the regulatory roles GmIFS2-mediated PPIs play across different complexes remain less characterized. Future proteomics study using soybean has the potential to profile different complexes co-existing in planta and further elucidate their metabolic roles.

Understanding the metabolic advantages of plant enzyme complexes offers novel strategies for PNP biomanufacturing. Although “mix-and-match” enzymes derived from different organisms based on their enzymatic activities has been the prevalent strategy to refactor efficient PNP pathways^45^, rebuilding pathways using interacting enzymes from the same host can have previously unknown metabolic advantages. In addition, while various artificial enzyme complexes have been constructed using scaffold DNA, RNA, or non-catalytic proteins^46^, CYP-mediated complexes can provide more structural and functional insights to build scaffold-free complexes with lower expression burden. In summary, yeast offers unparalleled advantages in the identification of PPIs and the biochemical characterization of enzyme complexes. The metabolic insights gained from yeast platforms will benefit PNPs manufacturing, as well as the study of plant metabolism and engineering.

## Materials and Methods

### Methods

#### Chemicals and reagents

Yeast nitrogen base (YNB) and amino acid mixtures were purchased from Sunrise Science Products. *P*-coumaric acid, formononetin and biochanin A was purchased from Sigma. Naringenin chalcone was purchased from MedCHemExpress. Naringenin was purchased from Alfa Aesar. Isoliquiritigenin was purchased from TCI. Liquiritigenin was purchased from CAYMAN Chemical Company. genistein, daidzein. Dextrose, yeast extract (YE), peptone, Luria-Bertani (LB) broth, LB agar, agar, acetonitrile, formic acid, and protease inhibitor (100x) were purchased from Thermo Fisher Scientific. All other chemicals, including antibiotics, were purchased from VWR International.

#### Microbes and culture conditions

*E. coli* Top 10, ccdB resistant *E. coli*, yeast CEN.PK2-1D and yeast strain THY.AP4 were used in this work. *E. coli* was used for plasmid construction: Top 10 for normal plasmids and the ccdB resistant strain for plasmids with ccdB gene. All *E. coli* strains harboring plasmids were cultivated in LB media or LB plates at 37℃ with 50 μg/mL of kanamycin or 100 μg/mL of carbenicillin as appropriate. Yeast CEN.PK2-1D was used for detecting PPI by BiFC or co-localization assay and studying biochemical functions. Yeast strain THY.AP4 was used for detecting PPI by split-ubiquitin Y2H. CEN.PK2-1D and THY.AP4 with plasmids were cultivated at 30°C, 400 rpm in appropriate synthetic drop-out (SD) liquid media or plates (0.17% YNB, 0.5% ammonium sulfate, 2% dextrose, and 370 amino acid drop-out mixture, and additional 2.5% agar for plates).

#### Plasmid construction for gene expression or for PPI identification in yeast

Reported isoflavonoids biosynthetic genes, *Gm4CL3, GmCHS7, GmCHI1B1, GmCHR5, GmIFS2, GmHID* from *G. max*, and *ATR1* from *A. thaliana* were codon-optimized and synthesized by Twist Bioscience. *GmIOMTs* were amplified from soybean cDNA. Gene sequences were listed in Supplementary Table 1. Oligonucleotide primers (Supplementary Table 2) were synthesized by Life Technologies. For gene expression in yeast, enzyme-encoding genes were inserted into pre-assembled Gateway-compatible plasmids that contain constitutive promoters and terminators to construct gene expression cassettes (Supplementary Table 3). For PPIs identification in yeast, enzyme-encoding genes with stop codon or without stop codon were inserted into pre-assembled Gateway-compatible plasmid that doesn’t contain constitutive promoters or terminators to construct entry plasmids (Supplementary Table 3). The gene expression cassettes were assembled into high-copy expression plasmids or low-copy plasmid using Gateway LR Clonase II Enzyme mix (Life Technologies), respectively. Plasmids were extracted using plasmid miniprep kits (Zymo Research), and the sequences were confirmed by Sanger Sequencing (Biotechnology Resource Center, Cornell University).

#### Yeast strain construction for isoflavonoids biosynthesis

Yeast strains (Supplementary Table 4) were constructed by yeast genome integrations or plasmid chemical transformation. To perform yeast genome integrations, DNA inserts containing pathway genes, auxotrophic marker genes, and 500- or 1000-bp genomic homologies, each harboring 25-bp overlaps between adjacent fragments, were PCR amplified from plasmids or yeast genomic DNA and transformed at equimolar ratios into yeast by electroporation at a voltage of 540 V, capacitance of 25 μF and infinite resistance with a Gene Pulser Xcell Total System electroporator (Bio-Rad). The transformed cells were immediately mixed with 1 mL of YPD medium and recovered at 30 °C, 400 rpm for 2 h, then harvested by centrifugation (8000 rpm, 1 min) and spread on selection plates for 2–3 days of growth. The correctly assembled constructs were verified through colony PCR analysis.

To perform the chemical transformation of plasmids into yeast host, chemically competent yeast cells were prepared using frozen-EZ yeast transformation II kits according to the manufacturer’s instructions. For single-plasmid transformation, 200 ng of plasmids were used while for double-plasmid transformation, 500 ng of each plasmid was used. After transformation, cells were spread on an appropriate selection plate and incubated at 30 °C for 2–3 days to allow for the growth of transformants.

#### Fermentation and sample preparation

Fresh yeast strains from plates were first incubated in 500 μL of SC, corresponding SD, or YP media supplemented with 2% dextrose as seed culture in deepwell plates at 30 °C, 400 rpm in replicates for 24 h and then transferred 50 μL seed culture into 450 μL of corresponding fresh media for 72 h. Certain amount of substrates (final concentration 2 mM of *p*-coumaric acid, 80 μM of naringenin chalcone or isoliquiritigenin, 125 μM of naringenin or liquiritigenin, 1 mg/L of genistein or daidzein) were fed for product production at the 24th hour. 150 μL supernatant of each culture was harvested analyzed by HPLC/ MS.

#### Metabolite analysis

Metabolites were identified and quantified by HPLC/Q-TOF (Agilent 1260 Infinity II/Agilent G6545B) in MS mode using positive ionization. For analysis, 1 μL of each culture sample was injected and separated in the ZORBAX RRHD Eclipse Plus C18 column (2.1 × 50 mm, 1.8 μm) (Agilent) with water with 0.1% formic acid (A) and acetonitrile with 0.1% formic acid (B) as the mobile phase. The gradient program was set as 0–1 min, 95% A; 1–11 min, 95%–5% A; 11–13 min, 5% A; 13–14 min, 5%–95% A; and 14–16 min, 95% A at a flow rate of 0.4 mL/min. The m/z value of the [M+H]^+^ adduct was then used to extract the ion chromatogram (with a mass error below 20 ppm) for compound identification by comparing the acquisition time and mass spectrum to that of the standard. Product concentrations were quantified by comparing the integrated peak area to that of the standard.

#### Characterization of PPIs by split-ubiquitin Y2H assay

The split-ubiquitin membrane Y2H system (termed here the SU system)^47^ was used to identify PPIs of GmIFS2 with other isoflavonoid biosynthetic enzymes. Genes encoding GmIFS2, GmCHS7, GmCHI1B1, GmCHR5 and GmHID and GmIOMTs without a stop codon were cloned from entry plasmids and recombined into pMetYC or pXN22 destination vectors by LR reaction to generate bait constructs and prey constructs, respectively. For the bait construct, the SUC2 peptide and the Cub-LexA-VP16 protein were added to the C-terminus of the GmIFS2, where SUC2 refers to the 19-residue signal peptide of Saccharomyces cerevisiae invertase and Cub-LexA-VP16 refers to the chimeric protein of the C-terminal half of ubiquitin (Cub) and a transcription factor cassette (LexA-VP16). For the prey constructs, GmCHS7, GmCHI1B1, GmCHR5 and GmHID and GmIOMTs were fused with the N-terminal half of a mutated ubiquitin (termed NubG) at their C-terminus (for example, GmCHS7-NubG). The bait and prey constructs were transformed into THY.AP4 yeast strains by electroporation and selected on synthetic dropout agar medium lacking leucine and tryptophan (SD−leu-trp agar plates). Triplicate colonies growing on SD-leu-trp plates were selected randomly and cultured in SD-leu-trp liquid medium overnight at 30 °C with shaking. The cell cultures were collected and five microliters each of 5-fold serial dilutions of this cell suspension were plated on both selective agar medium SD-leu-trp and detective agar medium SD-his. Images were taken after 4 days of growth at 30 °C.

#### Characterization of PPIs by BiFC assay

The BiFC assay was developed based on the split-mVenus fluorescent protein, which was split at the 210^th^ site, between the 10^th^ β-sheet and the 11^th^ β-sheet. The Addgene Yeast Gateway destination plasmids^48^ including pAG424/425-GPD-ccdB-eGFP or pAG424/425-GPD-eGFP-ccdB, were engineered by replacing the eGFP fragment with the NV and CV split fragments, respectively. Genes encoding *GmIFS2 (*without a stop codon), *GmCHS7*, *GmCHI1B1*, *GmCHR5*, *GmHID, GmIOMTs,* and negative control *Pm4’OMT* (synthesized by Twist Bioscience) (with or without a stop codon) were cloned from the entry plasmids and recombined into pAG425-GPD-CV-ccdB and the pAG424-GPD-ccdB-NV or pAG-424-GPD-NV-ccdB plasmids, respectively, using the Gateway LR Clonase II Enzyme mix (Life Technologies). Resultant plasmids were co-transformed into CEN.PK2-1D using the frozen-EZ yeast transformation II kits (Zymo Research). Corresponding yeast strains were cultivated at 30℃, 400 rpm in SD-trp-leu for three days. Cultured cells were washed with the PBS buffer (137 mM NaCl, 2.7 mM KCl, 10 mM Na_2_HPO_4_, and 1.8 mM KH_2_PO_4_) three times and resuspended with equ-volume PBS. Microscopic analysis was performed using a Zeiss LSM 710 Confocal Microscope (AxioObserver, with objective Plan-Apochromat 63X/1.40 Oil DIC M27) under the LineSequential scanning mode, with the excitation wavelength of 514 nm and signals of 519-620 nm. Transmitted light images (bright field and DIC) were also recorded.

#### Subcellular localization and co-localization analyses

To fuse the GmIFS2 and cytosolic isoflavonoid biosynthetic enzymes (GmCHS7, GmCHR5, GmCHI1B1 and GmHID) with eCFP and eGFP fluorescence proteins, the corresponding genes without a stop codon were cloned from the entry plasmids and were subsequently recombined into the pAG426-GPD-ccdB-eCFP and pAG424-GPD-ccdB-eGFP expression vectors from the *S. cerevisiae* Advanced Gateway Destination Vector Kit^48^ using LR Clonase II to generate the corresponding yeast expression constructs, respectively (Supplementary Table S3). Plasmids expressing GmIFS2-eCFP and GmCHS7 / GmCHIB1 / GmCHR5 / GmHID -eGFP were co-transformed into CEN.PK2-1D using the frozen-EZ yeast transformation II kits (Zymo Research). Resultant yeast strains were cultivated at 30 °C, 400 rpm in SD-trp-ura media for 16-18 hours and prepared as demonstrated above. Microscopic analysis was performed using a Zeiss LSM 710 Confocal Microscope (AxioObserver, with objective Plan-Apochromat 63X/1.40 Oil DIC M27). For the detection of eCFP fluorescence, the excitation wavelength was 405 nm, and signals of 442-510 nm were recorded by a PMT detector. For the detection of eGFP fluorescence, the excitation wavelength was 488 nm and signals of 491-588 nm were recorded by a PMT detector. Transmitted light images (bright field and DIC) were also recorded.

For characterization of the co-localization of a three-component complex, three fusion proteins, GmIFS2-DsRed, GmCHS7-eCFP and GmCHR5-eGFP were constructed by recombining gene fragments GmIFS2, GmCHS7, and GmCHR5 without a stop codon cloned from entry plasmids into the pAG423-GPD-ccdB-DsRed, pAG426-GPD-ccdB-eCFP and pAG424-GPD-ccdB-eGFP expression vectors, respectively. Plasmids expressing GmIFS2-DsRed, GmCHS7-eCFP and GmCHR5-eGFP were co-transformed into CEN.PK2-1D using the frozen-EZ yeast transformation II kits. Resultant yeast strains were cultivated and prepared as demonstrated above, and then performed microscopic analysis using a Zeiss LSM 710 Confocal Microscope. For the detection of DsRed fluorescence, the excitation wavelength was 561 nm, and signals of 563-706 nm were recorded by a PMT detector. For the detection of eCFP fluorescence, the excitation wavelength was 405 nm, and signals of 442-510 nm were recorded by a PMT detector. For the detection of eGFP fluorescence, the excitation wavelength was 488 nm and signals of 491-588 nm were recorded by a PMT detector. Transmitted light images (bright field and DIC) were also recorded.

#### Structure prediction and substrate docking using Alphafold2 and ChimeraX

All structure models are predicted by ChimeraX (v1.6) AlphaFold tool using Google Colab servers. Structures are visualized by ChimeraX. Amino residues within 5 angstroms to other proteins will be identified as the interface of the protein complex. Substrate docking was performed by Autodock.

#### Figure generation

Figures were generated through Prism v10 (Graphpad), BioRender.com, ZEISS Zen Lite, ChimeraX, Agilent Qualitative Analysis, and Microsoft Office 2016 (PowerPoint, Word, and Excel) whatever necessary.

## Contributions

C.L., J.H., and S.L. conceived and designed the experiments. C.L. and J.H. performed the experiments and analyzed the data. C.L., J.H., and S.L. wrote the paper.

## Competing Interests

The authors declare no competing interests.

## Acknowledgments

This work was supported by the National Institutes of Health National Institute on Deafness and Other Communication Disorders under award number R21DC019206 and National Center for Complementary and Integrative Health under award number R01AT012633, the National Science Foundation under Grants No. MCB-2338009, DBI-2019674, and IOS-2220733, and the Schwartz Research Fund Award. J. Han was supported by the Samuel C. Fleming Family Graduate Fellowship. We thank L. Li from USDA-ARS, Cornell University for kindly sharing the THY.AP4 yeast strains for Y2H assay. We thank the Imaging Facility (RRID: SCR_021741) of the Biotechnology Resource Center of Cornell Institute of Biotechnology. We thank F. Gong for providing gene templates during gene mining. We thank Y. Wu, A. Koganitsky, E. Parker Miller, and D. Han for their valuable comments on this manuscript.

## Extended Data

**Extended Data Figure 1.**
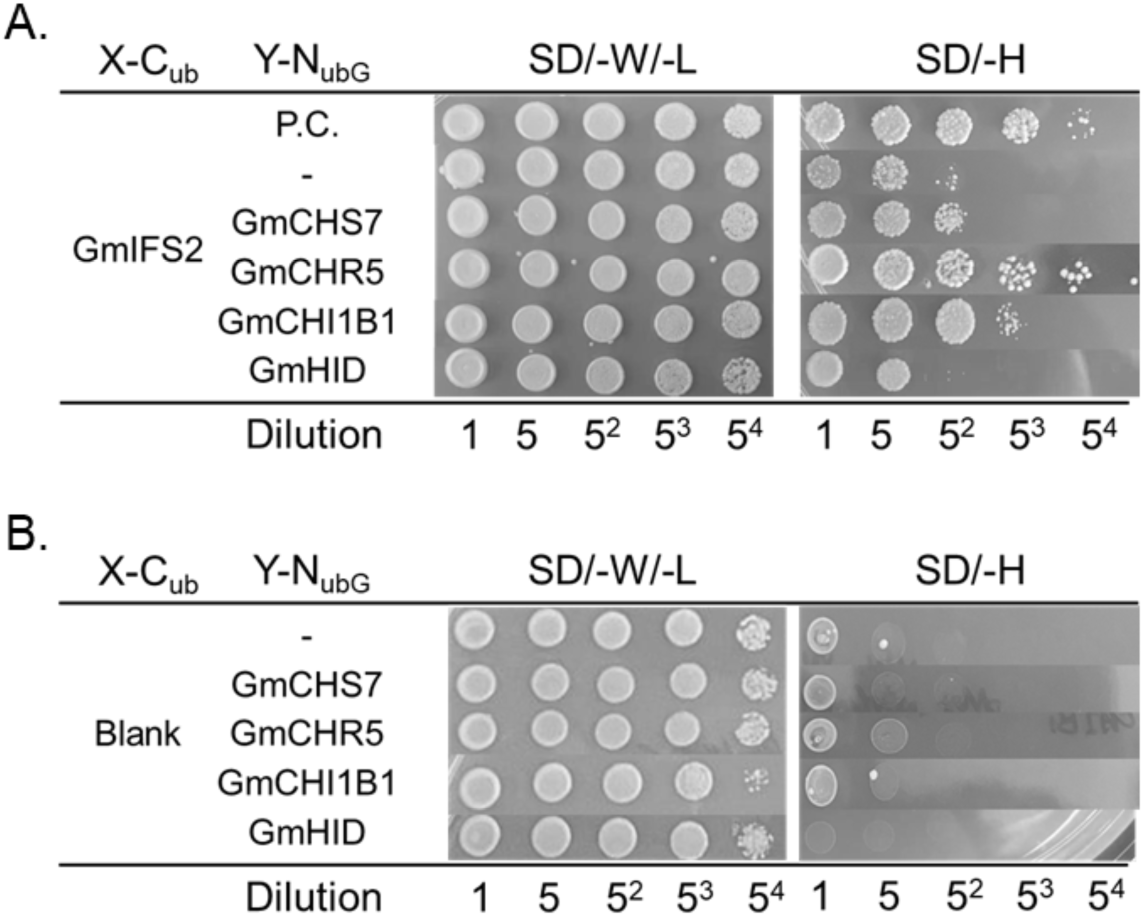
PPI detection of GmIFS2 with GmCHS7, GmCHR5, GmCHI1B1 and GmHID by split-ubiquitin (SU) Y2H assay. (A) Test groups. All the yeast cells co-expressed GmIFS2-C_ub_-LexA-VP16 and Y-N_ubG_, where Y refers to blank, GmCHS7, GmCHR5, GmCHI1B1 and GmHID. “Blank“ was used as negative control. P.C. refers to positive control. GmCHS7 homodimer was used as the positive control in this assay. **(B) Negative controls.** All the yeast cells co-expressed blank-C_ub_-LexA-VP16 and Y-N_ubG._ Figures are representatives of at least three biological replicates. Yeast cells were plated on SD-trp-leu plates (medium lacking tryptophan and leucine) for transformed strain selection and on SD-his plates (medium lacking histidine) for interaction detection.

**Extended Data Figure 2.**
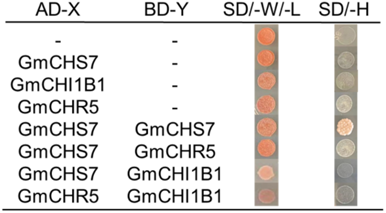
Detection of PPIs between GmCHS7, GmCHR5 and GmCHI1B1 by Gal4 Y2H assay. No interactions were identified in all the combinations besides GmCHS7 homodimer. PPIs between GmCHS7, GmCHR5 and GmCHI1B1 were also tested by BiFC assay, and no fluorescence signals were detected (data not shown).

**Extended Data Figure 3.**
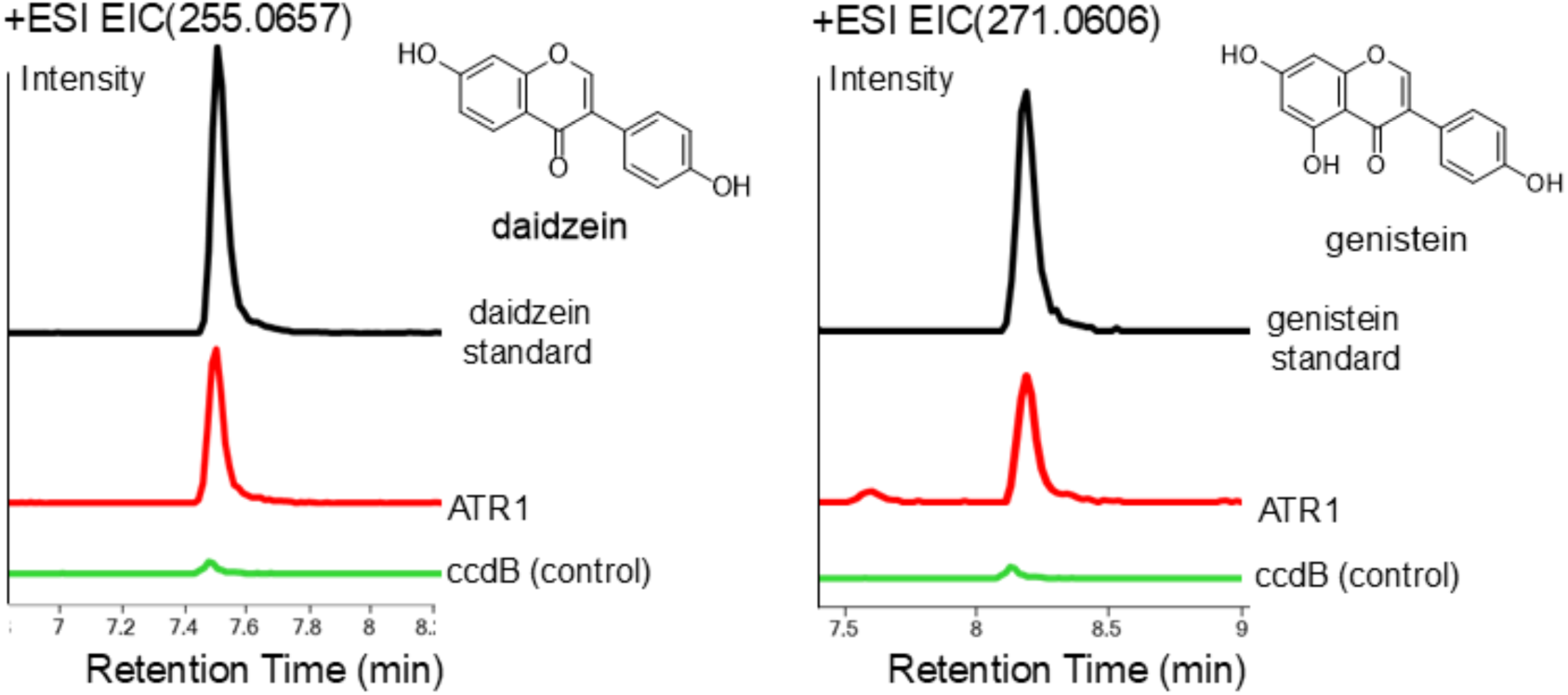
The metabolic activity of GmIFS2 in wild-type (without plant CYPR) yeast and ATR1 expressed yeast. GmIFS2 activity was characterized by monitoring the production of daidzein or genistein in yCL7. In wild type (without plant CYPR) yeast (yCL7 + ccdB), daidzein and genistein were barely produced. In contrast, in plant CYPR-efficient yeast (yCL7 + ATR1), daidzein and genistein were significantly produced. 2 mM of *p*-coumaric acid was fed and the supernatant was analyzed after 48 hours.

**Extended Data Figure 4.**
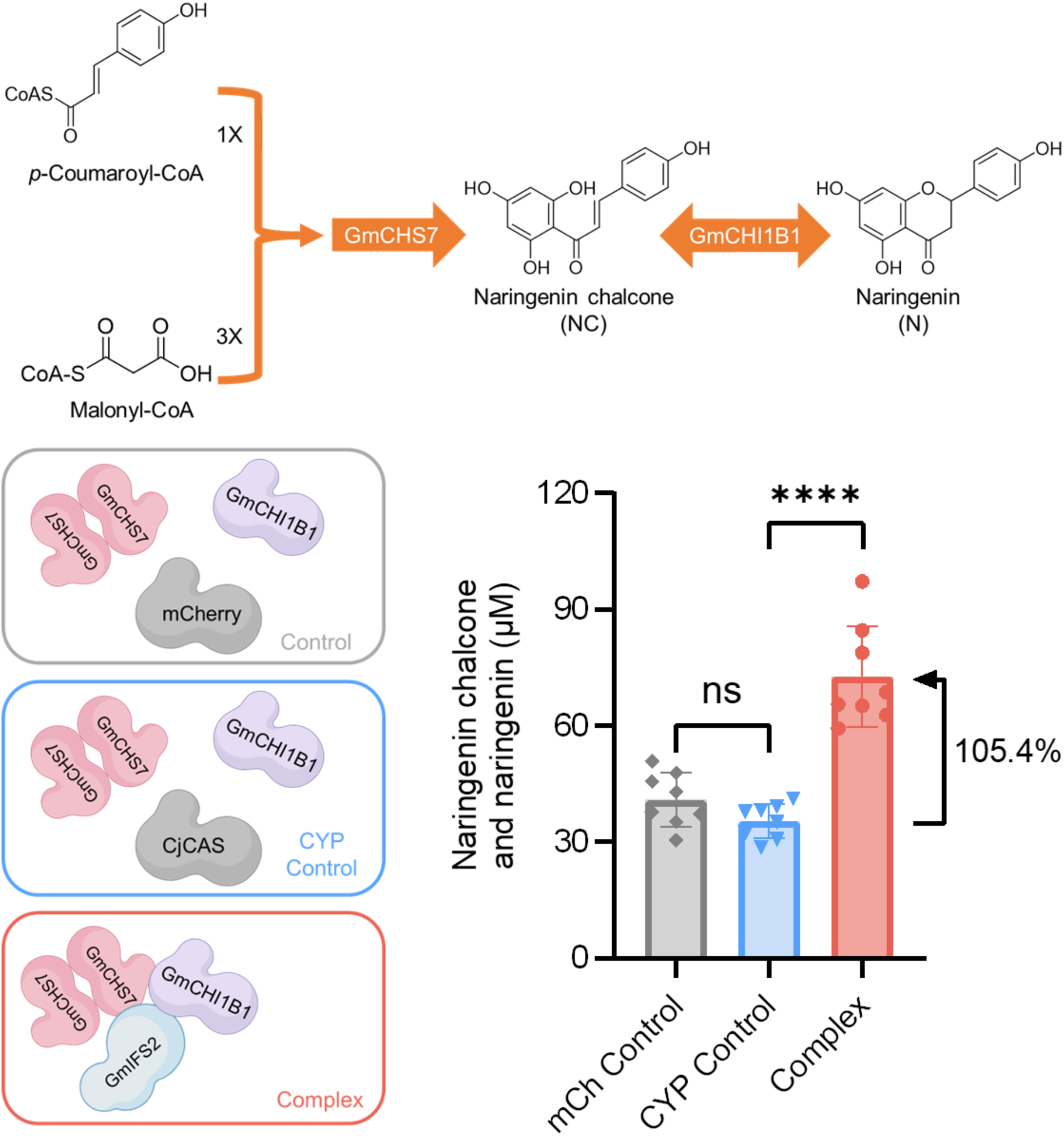
The metabolic activity of GmCHS7 when co-expressed with an unrelated CYP. Production of naringenin chalcone and naringenin by yeast strains harboring enzyme complexes (inactive GmIFS2 with GmCHS7) or mCh control (mCherry (mCh for short) with GmCHS7) and CYP control (a CYP, canadine synthase (CjCAS) from *Coptis Japonica*, with GmCHS7). 2 mM of p-coumaric acid was fed and the supernatant was analyzed after 48 hours. Data are means ± SD for at least six independent clones (t test, ns, not significant; ****P < 0.0001).

**Extended Data Figure 5.**
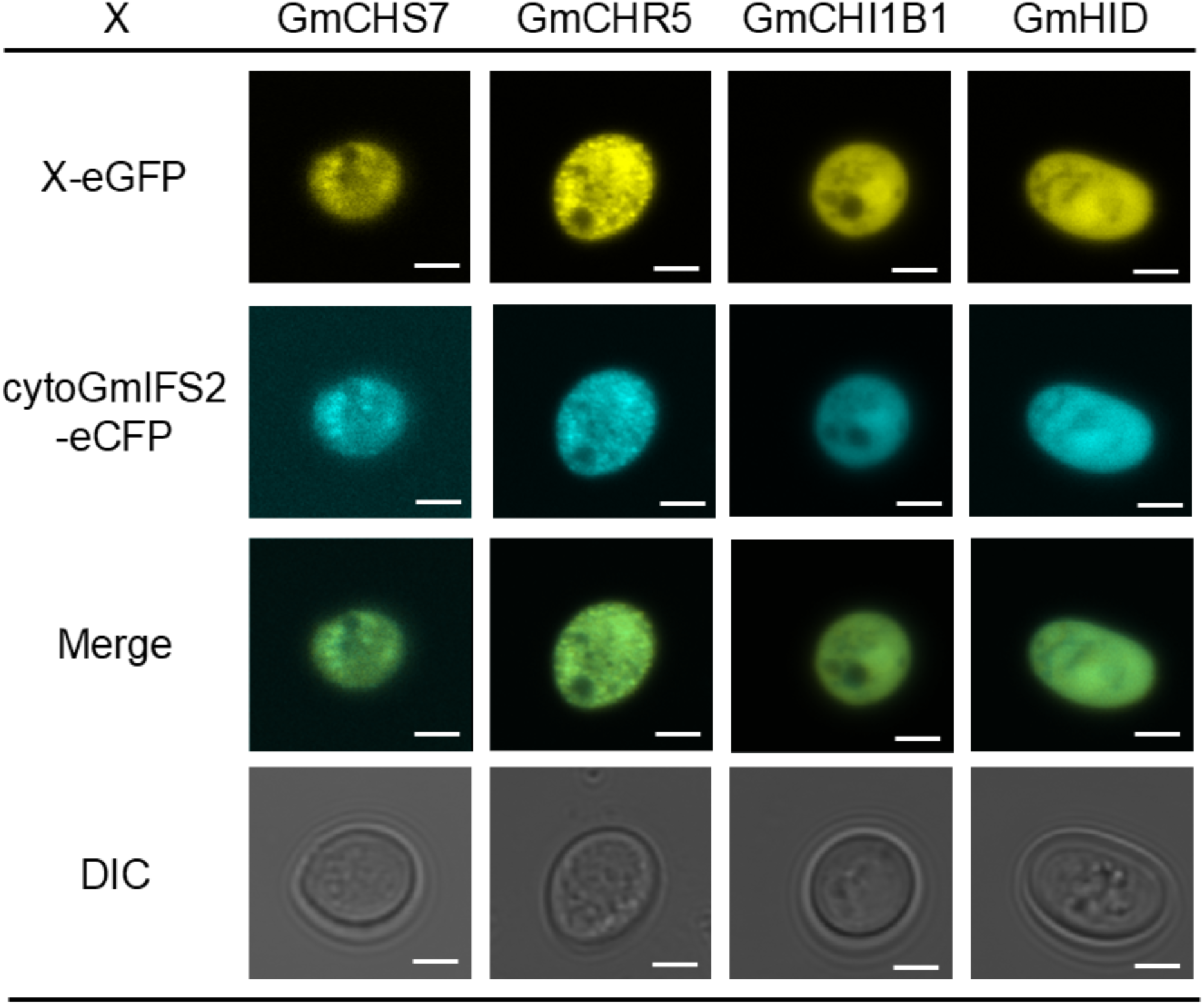
Localization assay of cytoGmIFS2 with GmCHS7, GmCHI1B1, GmCHR5 and GmHID in yeast cells. cytoGmIFS2 was fused with enhanced cyan fluorescent protein (eCFP) at the C-terminus. GmCHS7, GmCHB1, GmCHR5 and GmHID (negative control) were fused with eGFP at the C-terminus. All the enzymes were localized in cytosol. False colors were used for eCFP, eGFP and merged color. Cell morphology is observed with differential interference contrast (DIC). Scale bars = 2 μm.

**Extended Data Figure 6.**
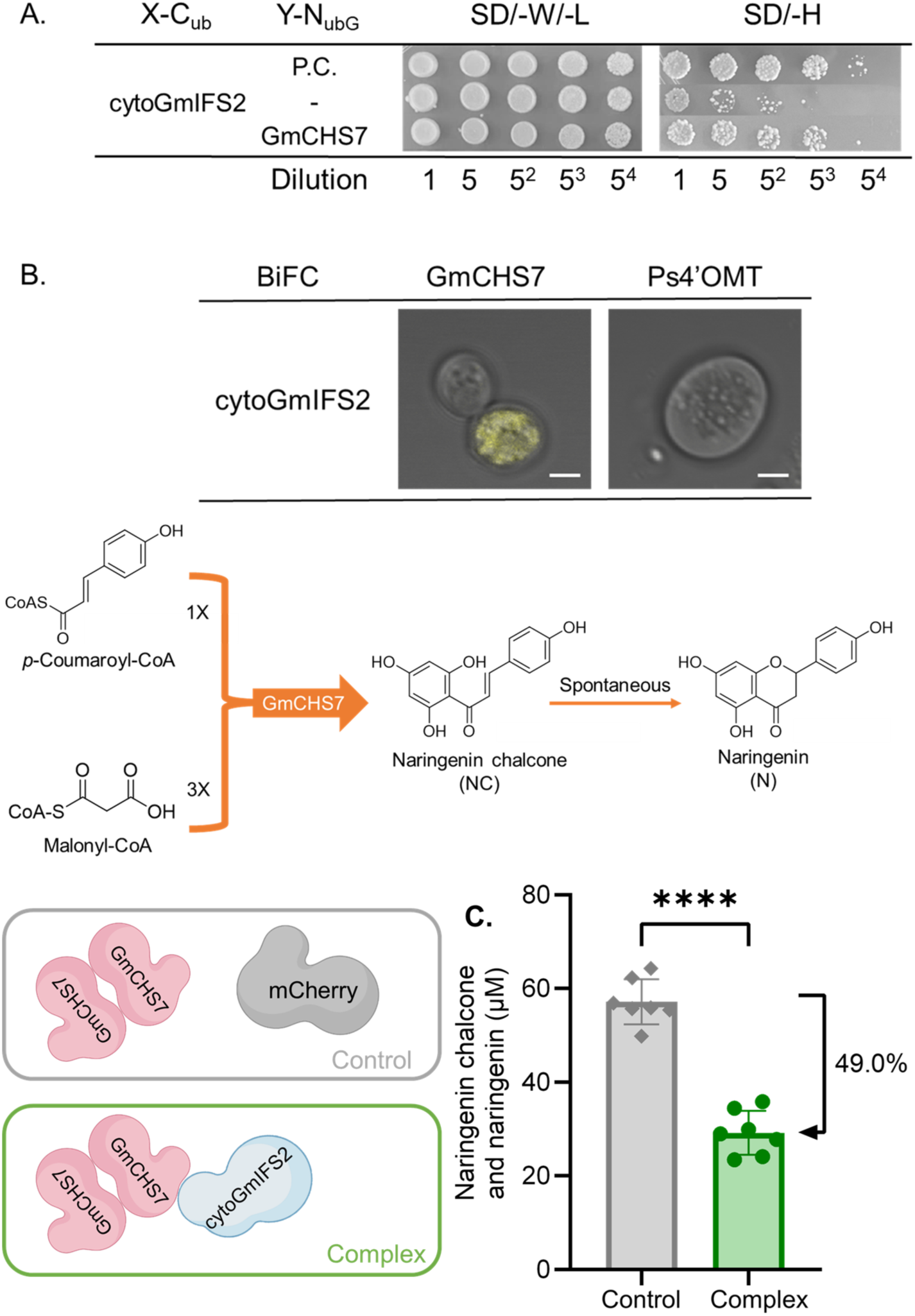
Metabolic effects of cytoGmIFS2 on the activity of GmCHS7 via PPI in yeast. **(A)** Y2H assay of cytoGmIFS2 and GmCHS7. **(B)** BiFC assay of cytoGmIFS2 and GmCHS7. **(C)** Production of naringenin chalcone and naringenin by yeast strains harboring enzyme complexes (inactive cytoGmIFS2 with GmCHS7) or control (mCherry with GmCHS7). 2 mM of p-coumaric acid was fed and the supernatant was analyzed after 48 hours. Data are means ± SD for at least six independent clones (t test, ****P < 0.0001).

**Extended Data Figure 7.**
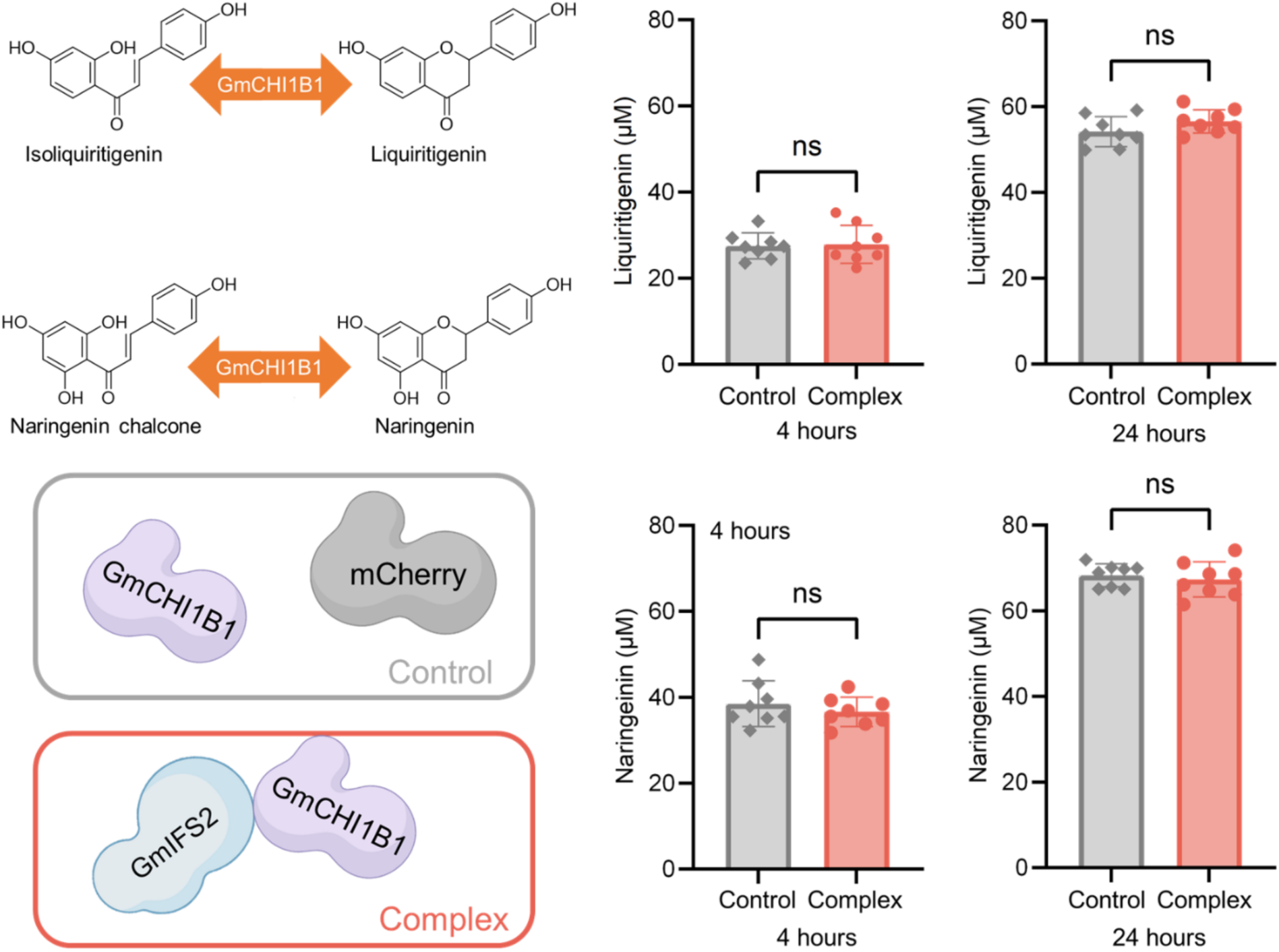
Production of liquiritigenin or naringenin by yeast strains harboring enzyme complexes (non-enzymatic GmIFS2 with GmCHI1B1) or control (mCherry with GmCHI1B1). 80 μM isoliquiritigenin or naringenin chalcone was fed and the supernatant was analyzed after 4 hours and 24 hours. Data are means ± SD for eight independent clones (t test, ns, not significant).

**Extended Data Figure 8.**
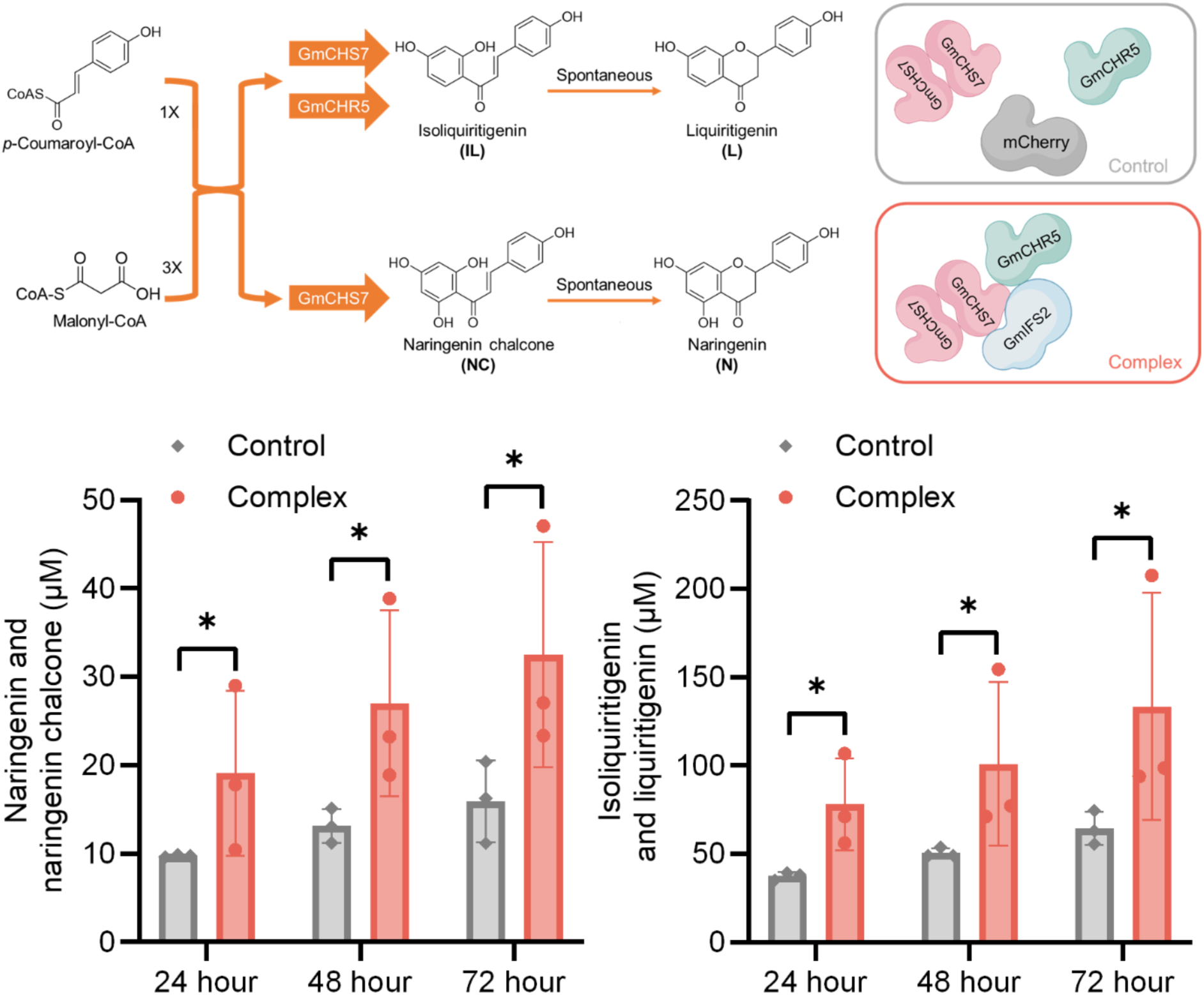
Production of flavonoids (naringenin chalcone and naringenin, NC+N) and deoxyflavonoids (isoliquiritigenin and liquiritigenin, IL+L) in yeast over 72 hours. 2 mM of *p*-coumaric acid was fed and the supernatant was analyzed after 24 hours, 48 hours and 72 hours. Data are means ± SD for three independent clones (t test, *P < 0.05).

**Extended Data Figure 9.**
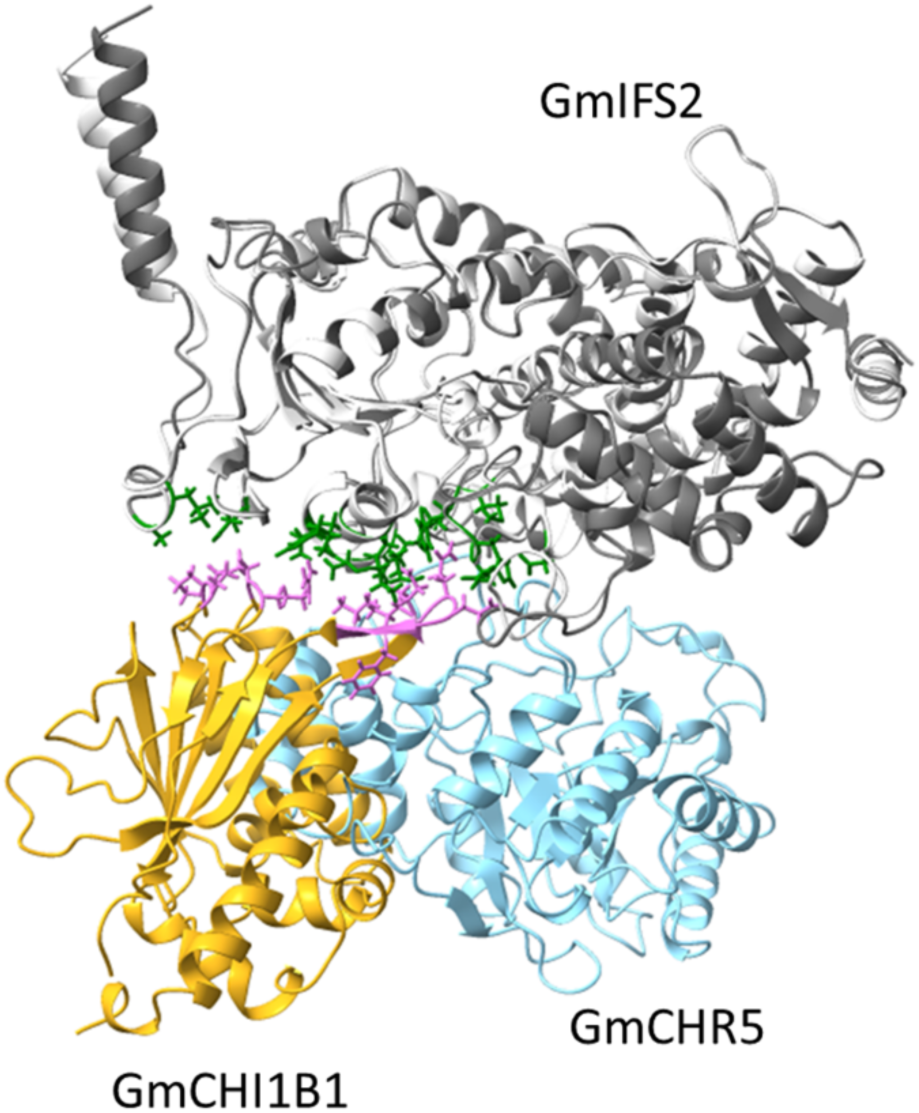
Interaction competition between GmCHI1B1 and GmCHR5 with GmIFS2 predicted by Alphafold-Multimer.

**Extended Data Figure 10.**
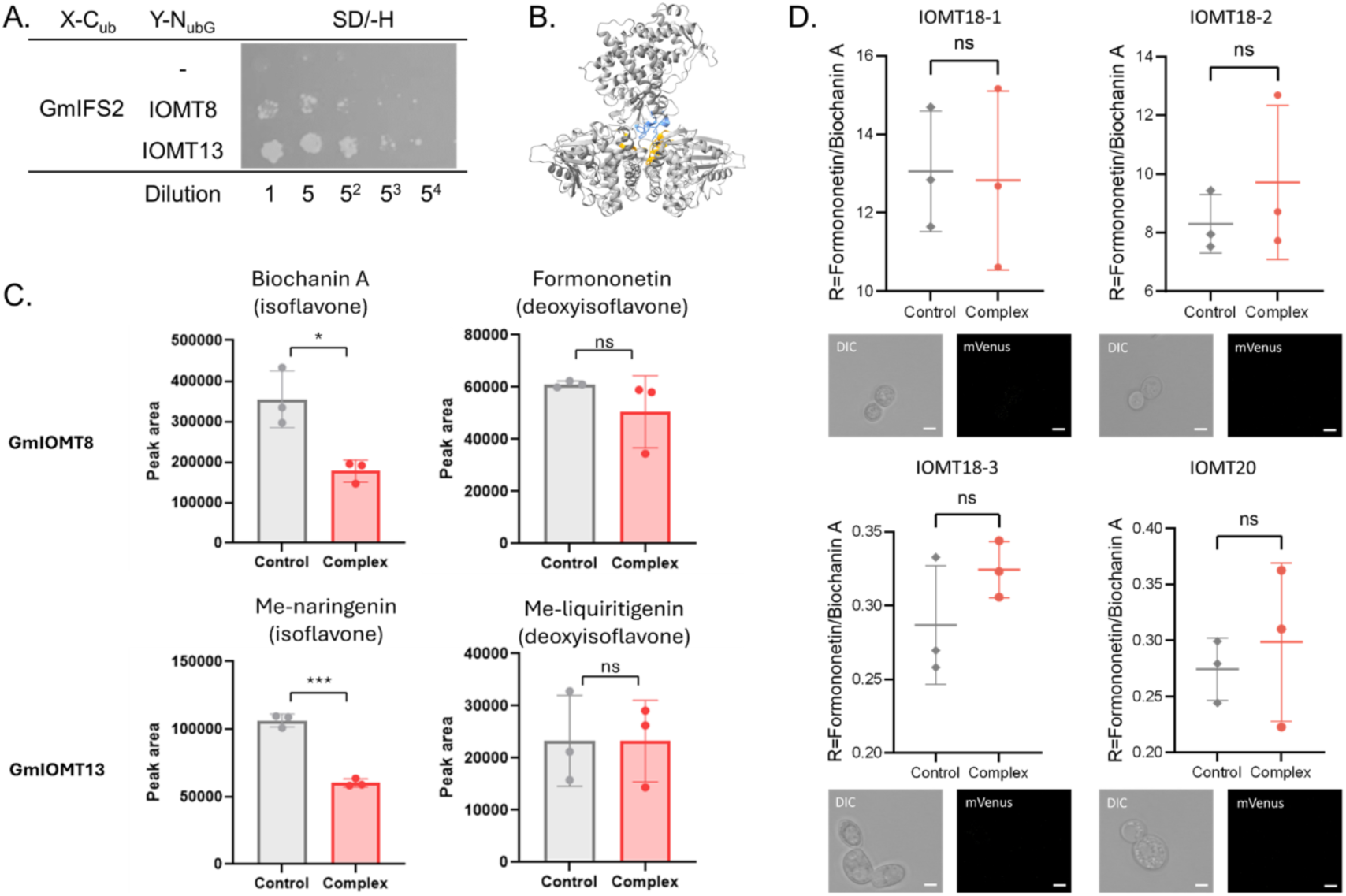
PPI and metabolic effects of GmIFS2 and GmIOMTs. **(A)**Y2H assay results for the interactions between GmIFS2 and IOMT8 or IOMT13. **(B)** The GmIFS2-IOMT8(dimer) enzyme complex structure predicted by Alphafold-Multimer. Amino residues within 5 Å to other proteins were identified as the interface of the protein complex. Interacting residues in GmIFS2 and GmIOMT8 were labeled in blue and yellow, respectively. **(C)** Production of GmIOMTs with or without IFS-mediated complex. 1 mg/L of substrate (genistein, daidzein, naringenin, and liquiritigenin) was fed and the supernatant was analyzed after 48 hours. Data are means ± SD for three independent clones (t test, *P < 0.05, ***P<0.001, ns, not significant). **(D)** BiFC assay results and R values for the interactions between GmIFS2 and non-interacting GmIOMTs. No fluorescence detected indicates no positive interactions were identified between these GmIOMTs with GmIFS2. R ratios were determined by dividing formononetin production by biochanin A production. Co-expressing non-interacting GmIOMTs with GmIFS2 didn’t redirect metabolic flux to deoxyflavonoids pathway. Data are means ± SD for three independent clones (t test, ns, not significant).

## Notes

### Competing Interest Statement

The authors have declared no competing interest.

